# Circuit mechanisms for chemical modulation of cortex-wide network interactions and exploration behavior

**DOI:** 10.1101/2020.06.25.171199

**Authors:** T. Pfeffer, A. Ponce-Alvarez, T. Meindertsma, C. Gahnström, R. L. van den Brink, G. Nolte, K. Tsetsos, A.K. Engel, G. Deco, T.H. Donner

## Abstract

Influential accounts postulate distinct roles of the catecholamine and acetylcholine neuromodulatory systems in cognition and behavior. But previous work found similar effects of these modulators on the response properties of individual cortical neurons. Here, we report a double dissociation between catecholamine and acetylcholine effects at the level of cortex-wide network interactions in humans. A pharmacological boost of catecholamine levels increased cortex-wide interactions during a visual task, but not rest. Conversely, an acetylcholine-boost decreased correlations during rest, but not task. Cortical circuit modeling explained this dissociation by differential changes in two circuit properties: the local excitation-inhibition balance (more strongly altered by catecholamines) and intracortical transmission (more strongly reduced by acetylcholine). The inferred catecholaminergic mechanism also predicted increased behavioral exploration, which we confirmed in human behavior during both a perceptual and value-based choice task. In sum, we identified specific circuit mechanisms for shaping cortex-wide network interactions and behavior by key neuromodulatory systems.

## Introduction

The catecholaminergic (noradrenergic and dopaminergic) and cholinergic modulatory systems of the brainstem are important regulators of global brain state and cognition (Arnsten, 2015; Aston-Jones and Cohen, 2005; Bear and Singer, 1986; Cools, 2019; Harris and Thiele, 2011; Robbins and Arnsten, 2009). Their brainstem centers send ascending projections to large parts of the cerebral cortex (Aston-Jones and Cohen, 2005; Breton-Provencher and Sur, 2019; Schwarz and Luo, 2015), which is equipped with similarly widely distributed receptors for those neuromodulators (van den Brink et al., 2019; Burt et al., 2018). Consequently, these systems are in an ideal position to shape cortex-wide network activity in a coordinated fashion. Indeed, mounting evidence indicates that these systems have a profound impact on large-scale correlations in cortical activity, as measured by neuroimaging or electrophysiological mass signals (van den Brink et al., 2016, 2018, 2019; Coull et al., 1999; Leopold et al., 2003; Turchi et al., 2018).

Influential theoretical accounts postulate highly specific roles of the catecholaminergic and cholinergic systems in the regulation of cognition and behavior (Aston-Jones and Cohen, 2005; Montague et al., 2004; Yu and Dayan, 2005). One prominent idea holds that catecholamines increase the responsivity (‘gain’) of neuronal populations to synaptic input (Aston-Jones and Cohen, 2005; Eldar et al., 2013; Servan-Schreiber et al., 1990). Through this mechanism, catecholamines can increase behavioral variability to promote exploratory decision-making when required by the environmental context (e.g. to learn about new sources of reward) (Aston-Jones and Cohen, 2005). Acetylcholine, on the other hand, has been proposed to reduce the impact of prior knowledge (intra-cortical signaling) relative to new information (bottom-up signaling) (Yu and Dayan, 2005). Such specific functional roles imply that these modulators should also have specific effects on the activity of the cortical circuits that implement cognitive computation.

Due to a lack of experimental comparisons, physiological evidence for a distinct shaping of cortical activity and behavior through catecholamines and acetylcholine is sparse. At the cellular level, catecholamines and acetylcholine, in fact, both increase the gain of cortical neurons (Disney et al., 2007; Herrero et al., 2008; Hurley et al., 2004; Polack et al., 2013), which translates into an increased neuronal ‘signal-to-noise ratio’(Aston-Jones and Cohen, 2005; Robbins and Arnsten, 2009). However, the relative magnitudes of the catecholaminergic versus cholinergic gain modulations have not been assessed. Further, while some studies have shown a suppression of intra-cortical signaling through acetylcholine (Hsieh et al., 2000; Roberts et al., 2005; Silver et al., 2008), it remains unknown whether the same holds for catecholamines, to a similar degree. A direct comparison between the circuit-level, large-scale, and behavioral effects of catecholamines and acetylcholine is required for pinpointing potential differences between the modulatory systems, which might emerge at any of the above levels.

Here, we set out to conduct such a direct comparison. Our approach was inspired by insights from theoretical neuroscience (Deco et al., 2014; Shine et al., 2018) – specifically, that (i) the large-scale interaction of relatively subtle local microcircuit effects can give rise to substantial effects at the level of cortex-wide network dynamics, and (ii) just as for single neurons (Servan-Schreiber et al., 1990), the network effects of gain modulation should depend on the external drive the network receives. We performed a direct comparison between the effects of placebo-controlled pharmacological increases of catecholamine or acetylcholine levels on large-scale cortical interactions in humans in two behavioral contexts: a visual task (i.e., external drive) and rest (absence of drive). This yielded an unexpected, context-dependent double-dissociation between the effects of catecholamines and acetylcholine. We then used computational modeling, across multiple levels of cortical organization, to infer the circuit mechanisms underlying this dissociation. Model simulations explained the catecholaminergic effect through a net increase in the population gain of local cortical regions, likely mediated by a ‘disinhibition’ of the underlying microcircuits. By contrast, model simulations explained the cholinergic effect through a suppression of intra-cortical signal transmission combined with weaker net gain modulation. The catecholaminergic circuit disinhibition also predicted an increase in behavioral choice variability in a circuit model for decision-making. We confirmed this prediction for human behavior in two datasets under the same manipulation of catecholamine levels, for the domains of perceptual and value-based decision-making (Sugrue et al., 2004, 2005). Our results provide critical constraints for future computational theories of neuromodulatory function and set the stage for the development of non-invasive biomarkers for the integrity of neuromodulatory function.

## Results

We increased central catecholamine and acetylcholine levels through the placebo-controlled administration of atomoxetine and donepezil, respectively (Fig. 1A, left; see Methods; data re-analyzed from a previous report on local cortical variability (Pfeffer et al., 2018)). Atomoxetine is a selective noradrenaline reuptake inhibitor. Consequently, atomoxetine increases noradrenaline levels across cortex (Robbins and Arnsten, 2009) and dopamine levels in its more restricted cortical projection targets (mainly frontal cortex) (Bymaster et al., 2002). Donepezil is a cholinesterase inhibitor (Silver et al., 2008), which blocks the enzymatic breakdown of synaptic acetylcholine and thus boosts cortical acetylcholine levels. Both drugs are routinely used in the clinical practice for treating important neuropsychiatric disorders, such as attention deficit hyperactivity disorder (atomoxetine) and Alzheimer’s disease (donepezil).

**Fig. 1.**
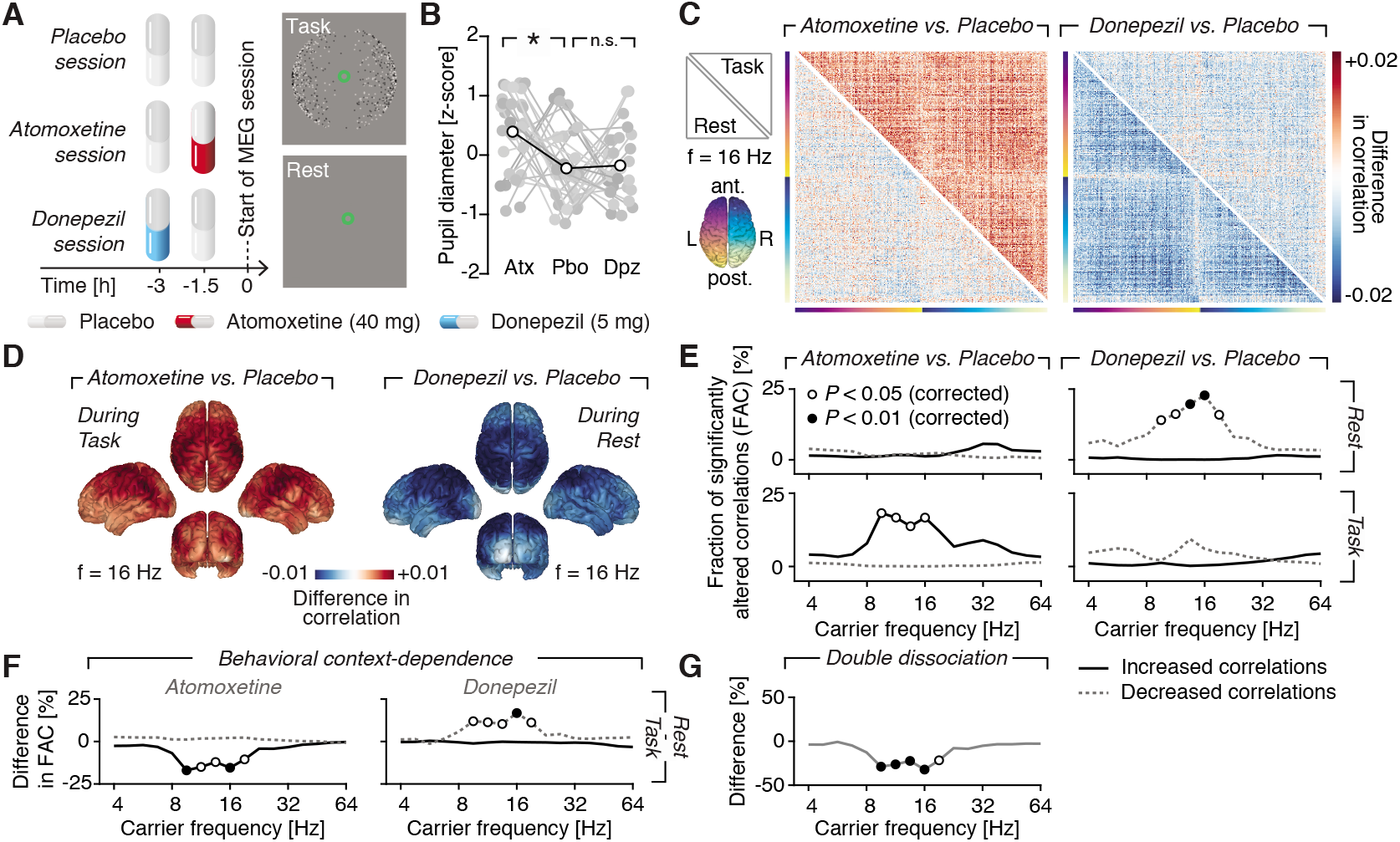
Dissociated catecholaminergic and cholinergic effects on cortex-wide correlations in activity. **(A)** Experimental design. Top: Atomoxetine (40 mg), donepezil (5 mg), or a visually indistinguishable placebo was administered before each session. Bottom: MEG activity was recorded during a visual task (left) or eyes-open ‘rest’ (right). **(B)** Drug effect on baseline pupil diameter (rest and task collapsed. Atx, atomoxetine; Pbo, placebo; Dpz, donepezil; * *P* < 0.05, paired two-sided permutation test) **(C)** Drug effects on cortex-wide activity correlations (at 16 Hz), for task (upper triangle) and rest (lower triangle). Left: atomoxetine – placebo; Right: donepezil – placebo. **(D)** Cortical distribution of drug effects on correlations. Left: Atomoxetine – placebo (task). Right: Donepezil – placebo (rest). **(E)** Frequency spectrum of drug effects on the fraction of significantly (*P*<0.05, paired t-test) altered correlations across brain regions, for atomoxetine (left) and donepezil (left) as well as for rest (top) and task (bottom). Fractions of significantly increased (solid black lines) and decreased (dashed gray lines) correlations are shown separately. **(F)** Effect of behavioral context on correlations (difference between upper and lower rows in E). **(G)** Spectrum of double dissociation between atomoxetine and donepezil effects, measured as the difference between (panel E): solid black line (left) and dashed gray line (right). Open circles, *P* < 0.05; filled circles, *P* < 0.01 (paired two-sided single-threshold permutation test).

Atomoxetine (catecholamines), but not donepezil (acetylcholine), increased pupil size (Fig. 1B, Fig. S1, an established peripheral marker of central arousal state (Breton-Provencher and Sur, 2019; de Gee et al., 2017; Joshi et al., 2016; McGinley et al., 2015; Reimer et al., 2016).

The rationale of our analyses, and hence organization of the Results is as follows. We (i) measured the effects of these pharmacological interventions on large-scale cortical network dynamics (assessed with magnetoencephalography; MEG) and behavior, and (ii) simulated cortical circuit models in order to develop a mechanistic understanding of our empirical results. By (iii) extending the model into a circuit that generates selection (i.e., choice) behavior, we turned the mechanistic inference into a prediction for behavior, which (iv) we finally confirmed in two independent datasets that probed into both, perceptual and value-based decision-making (Sugrue et al., 2005).

### Distinct, context-dependent drug effects on large-scale network dynamics

Large-scale cortical network interactions were quantified as the frequency-resolved, cortex-wide correlations of intrinsic activity fluctuations (Fig. 1C). Critically, we measured these intrinsic correlations in two behavioral contexts: a visual task with continuous input and eyes-open ‘rest’ (Fig. 1A, right). The task entailed the continuous presentation of an ambiguous visual stimulus, which, in turn, induced spontaneous and ongoing alternations in perception (Leopold and Logothetis, 1999) (Fig. 1A and Movie S1). During MEG blocks, we asked participants to silently count perceived perceptual alternations and report the total count at the end of each run. Consequently, we could assess intrinsic fluctuations in MEG activity in the absence of transients in visual input and motor movements. In separate blocks, participants reported each perceived alternation with an immediate button press.

We used a previously established approach (Hipp et al., 2012) that attenuates spurious correlations due to signal leakage (see Methods and Fig. S2A for illustration). We computed pairwise correlations between 400 cortical locations and compiled them into a matrix, separately for a range of carrier frequencies. Averaged across the placebo-rest condition, this yielded a similar spatial and spectral structure of correlations as previously reported (Hipp et al., 2012) (Fig. S2B,C). We then compared the correlation matrices between task and rest (Fig. S3), and between each drug condition and placebo condition (Fig. S4). Neuromodulators may potentially cause correlations between cortical mass signals to shift in a common direction (e.g., toward larger positive correlations), or change in magnitude (e.g. shift toward more negative and more positive correlations (Eldar et al., 2013)), depending on the underlying mechanism (van den Brink et al., 2019). To statistically assess the differences between our experimental condition in an unbiased fashion, we computed the fraction of significantly increased and decreased correlations, separately for each frequency bin. We then tested those fractions for their deviation from the expected chance-level, while accounting for multiple comparisons across frequencies (Methods and Fig. S3, S4).

Atomoxetine increased correlations across most pairs of regions during task (Fig. 1C, left; upper triangular part; Fig. S4A). This effect was evident in all four cerebral lobes (Fig. 1D, left) and peaked in the ‘alpha/beta’ frequency band (9.51-16 Hz; Fig. 1E, left). The effect was absent during rest (Fig. 1E; upper vs. lower triangular part in Fig. 1C).

In sharp contrast, donepezil (acetylcholine) decreased correlations across most region pairs, but only during rest (Fig. 1C, right, lower triangular part; Fig. 1E, right; Fig. S4B). Consequently, both drugs had opposite effects on correlations, dependent on behavioral context within overlapping frequency bands (Fig. 1F). These opposite effects translated into a robust, frequency-specific and context-dependent double dissociation between the atomoxetine and donepezil effects on cortical network dynamics (Fig. 1G; all *P*-values < 0.01 for the range: 9.51-16 Hz).

The double dissociation was neither present at the level of local activity fluctuations (see Pfeffer et al., 2018 and Fig. S5), nor did it depend on specific choices of analysis parameters (Fig. S6-7).

### Distinct changes in circuit parameters explain drug effects on large-scale network dynamics

Catecholamines and acetylcholine both increase the gain of cortical neurons (Aston-Jones and Cohen, 2005; Herrero et al., 2008; Polack et al., 2013; Servan-Schreiber et al., 1990). How, then, did the dissociation between their large-scale effects arise? To illuminate this question, we modeled the mass activity of coupled cortical regions (‘nodes’), each of which was composed of an interconnected excitatory and inhibitory neural population (Wilson and Cowan, 1972) (Fig. 2A, B, see also Methods and Supplementary Discussion). The model had four free parameters: the background inputs to excitatory (*b_E_*) and inhibitory (*b_I_*,) populations, the slope of the input-output function (‘gain’ at the neural population level) and a global coupling parameter.

**Fig. 2.**
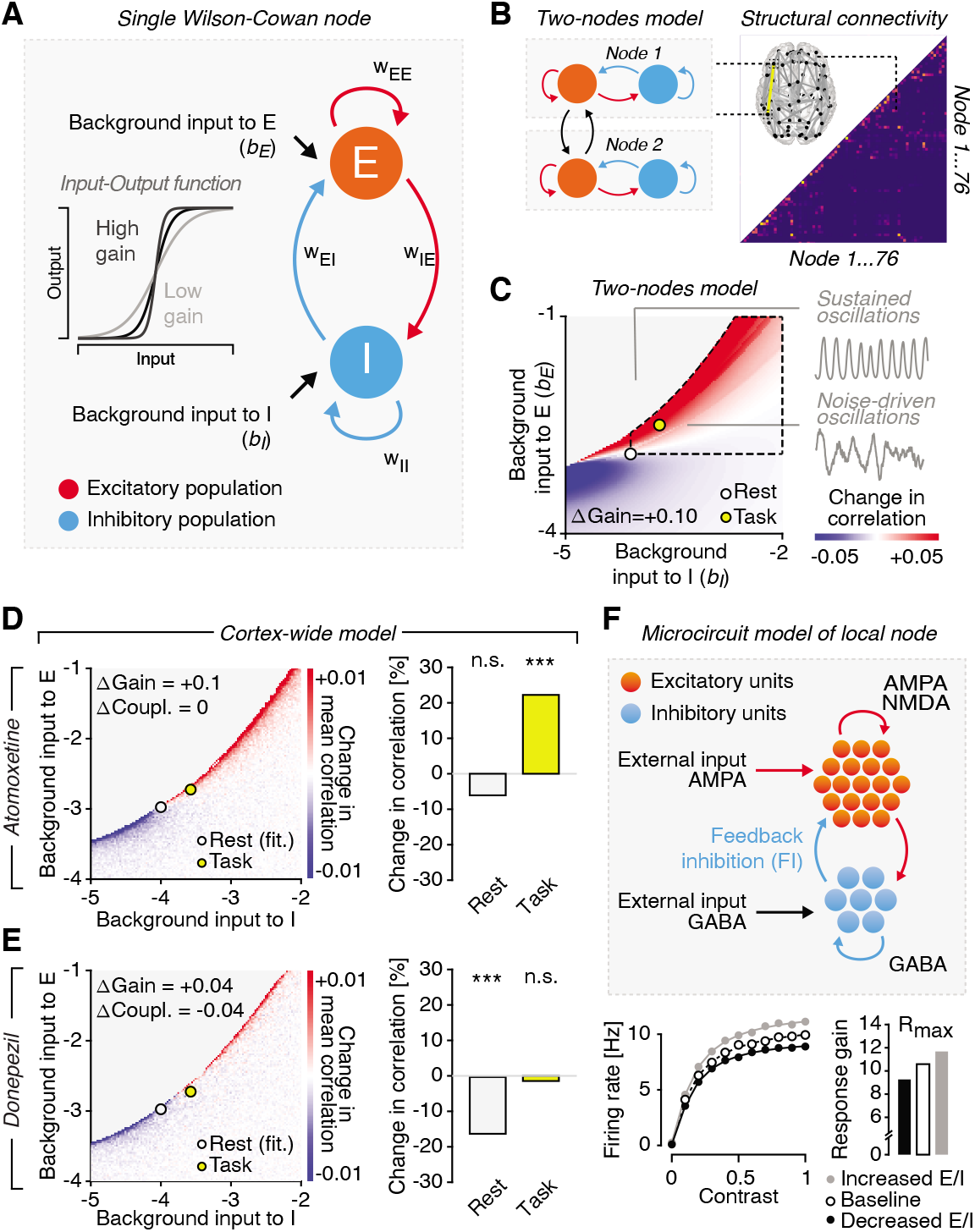
Circuit mechanisms of context-dependent effects on cortex-wide correlations. **(A)** Schematic of a single node (brain region) consisting of an excitatory (E) and an inhibitory population (I), with full connectivity and independent background input to E and I (Wilson-Cowan model(Wilson and Cowan, 1972)). Inset, input-output function of each population for various gain parameters (slope of the input-output function). **(B)** Left: Correlations were computed between the firing rates of the E populations of two or more nodes. Right: For the cortex-wide model, an estimate of the human structural connectome was used to connect a total of 76 nodes, and the model was fitted to the rest-placebo data (see Methods). **(C)** Change in correlation under an increase in gain (+0.1) in the (*b_E_, b_I_*)-plane of the ‘two-nodes model’. Inset: sustained and noise-driven oscillations. The area defined by the dashed black line highlights the assumed task-related shift in the (*b_E_, b_I_*)-plane. **(D)** Effect of gain increase (+0.1) across all 76×76 node pairs (right; white circle: rest; yellow circle: task) **(E)** As D, but for donepezil with gain increase by +0.04 (see Fig. S9 for +0.1) and decrease in global coupling (−0.04). **(F)** Top: Architecture of microcircuit consisting of excitatory and inhibitory integrate-and-fire neurons (all-to-all connectivity). Bottom, left: Effect of change in E/I (feedback inhibition) on gain for increases (decreased E/I; black line; filled black circles) and decreases in feedback inhibition (increased E/I; gray line; filled gray circles), with respect to baseline (dashed line; open circles). Bottom, right: fitted response gain parameter (*R_max_*) of the stimulus-response function for three levels of E/I.

Our simulations were constrained by two assumptions derived from established physiology (see also Supplementary Discussion). First, we assumed that cortical mass activity exhibits noise-driven oscillations, as opposed to sustained oscillations. In the noise-driven (also referred to as fluctuation-driven) regime, stochastic fluctuations in activity drive damped oscillations in the local nodes. Superposition of such damped oscillations, triggered at random moments in time, give rise to the ongoing variations in the amplitude of band-limited activity. Second, we assumed increased background input (*b*) to excitatory populations (*b_E_*) and inhibitory populations (*b_I_*,) in many cortical regions during task (Haider et al., 2013). This assumption rests on the notion that our visual task increased the input to visual as well as higher-order ‘task-related’ cortical regions, and affected both excitatory and inhibitory neural populations in these regions. Indeed, sensory input increases not only excitation (i.e., drive of pyramidal cells), but also inhibition (i.e., drive of interneurons) in sensory cortex (McCasland and Hibbard, 1997; Swadlow, 2002). Correspondingly, in the model, task increased the background input to excitatory as well as inhibitory populations relative to rest, resulting in an upward-rightward shift in the (*b_E_, b_I_*)-plane. Fig. 2C illustrates these two assumptions in terms of the gray shaded area (top left; sustained oscillations) and the dashed black outline (task-related increase in background input; shown for a version of the model made up of only two nodes).

We used the model to test the hypothesis that differences in gain modulation and/or changes in global coupling could explain the double dissociation observed in the data. Gain increase under catecholamines has been established for the input-output functions of single cortical neurons (Aston-Jones and Cohen, 2005; Hurley et al., 2004; Servan-Schreiber et al., 1990) as well as of neural mass activity assessed with neuroimaging (Eldar et al., 2013; Shine et al., 2018). Our simulations showed that an increase in gain was, in fact, sufficient to explain the context-dependent effect of atomoxetine on cortical correlations (Fig. 2C). Just as observed in the empirical data (Fig. 1E, left), increasing the gain in the ‘two-node model’ produced distinct changes in correlations for different contexts situated in the (*b_E_*, *b_I_*)-plane (i.e., different levels of background drive; rest: white circle; task: yellow circle; Fig. 2C). In a realistic model of the whole cortex (Fig. 2B, right), which was fitted to the measured correlation matrix for rest-placebo (Fig. S8F), an increase in gain boosted correlations in the same context-dependent fashion, with no change at rest (Fig. 2D, light gray circle or bar), but a robust increase during task (Fig. 2D, yellow circle or bar).

Increases in neural gain result from complex synaptic interactions (Ferguson and Cardin, 2020), at a spatial scale smaller than the one of our neural mass model of entire cortical nodes. We reasoned that the inferred catecholaminergic gain increase may have resulted from a catecholaminergic increase the ratio between excitation and inhibition (henceforth termed ‘E/I’) within cortical microcircuits (Froemke, 2015; Murphy and Miller, 2003; Polack et al., 2013). Indeed, noradrenaline tonically suppresses ongoing inhibitory inputs to pyramidal cells (Martins and Froemke, 2015; Polack et al., 2013), which may translate into an increase in the gain of the whole microcircuit. We simulated cortical microcircuit model to test this idea (Methods; Fig. 2F, top; Fig. S10). The microcircuit model was made up of recurrently connected excitatory and inhibitory conductance-based spiking neurons. We increased the circuit’s E/I by decreasing the strength of feedback inhibition and of quantified the effect on response gain the input-output function of the excitatory cells of the circuit (Methods). In line with our reasoning, increasing E/I translated into a response gain increase (Fig. 2F, bottom).

Our neural mass model could also explain the opposite, context-dependent effect of acetylcholine on large-scale cortical dynamics (Fig. 1E, right). While acetylcholine, like catecholamines, increases the gain of single neurons (Disney et al., 2007; Herrero et al., 2008, 2017; Soma et al., 2012), cholinergic and noradrenergic effects on E/I (i.e., gain) differ: the cholinergic E/I (i.e., gain) increase affects a smaller fraction of neurons in the circuit for a shorter duration (Froemke, 2015; Froemke et al., 2007), which likely translates into a smaller impact on the microcircuit’s (i.e. node’s) net gain (see Discussion). An increase in gain, indeed smaller than the catecholaminergic increase, selectively decreased correlations during rest (not task) – but, critically, this gain modulation had to be combined with a decrease in the model’s global coupling parameter (Fig. 2E). Such a decreased global coupling is in line with a reduction of intracortical (lateral and/or feedback) signaling observed in sensory cortex (Hsieh et al., 2000; Roberts et al., 2005; Silver et al., 2008) (see Discussion).

### E/I increase under catecholamines also accounts for increased behavioral variability

The above circuit modeling insights, specifically the cortical E/I increase under catecholamines, also accounted for the observed drug effects on visually-guided behavior, further validating our conclusions (Fig. 3). Atomoxetine (not donepezil) increased the number of perceptual alternations reported by participants during MEG (Fig. 3A), a simple readout of behavioral variability (Renart and Machens, 2014). This effect was not due to a change in eye movements or blinks (Pfeffer et al., 2018), and it was evident both when participants silently counted the perceptual transitions and when they reported each perceptual transition with an immediate button press (Fig. S11).

**Fig. 3.**
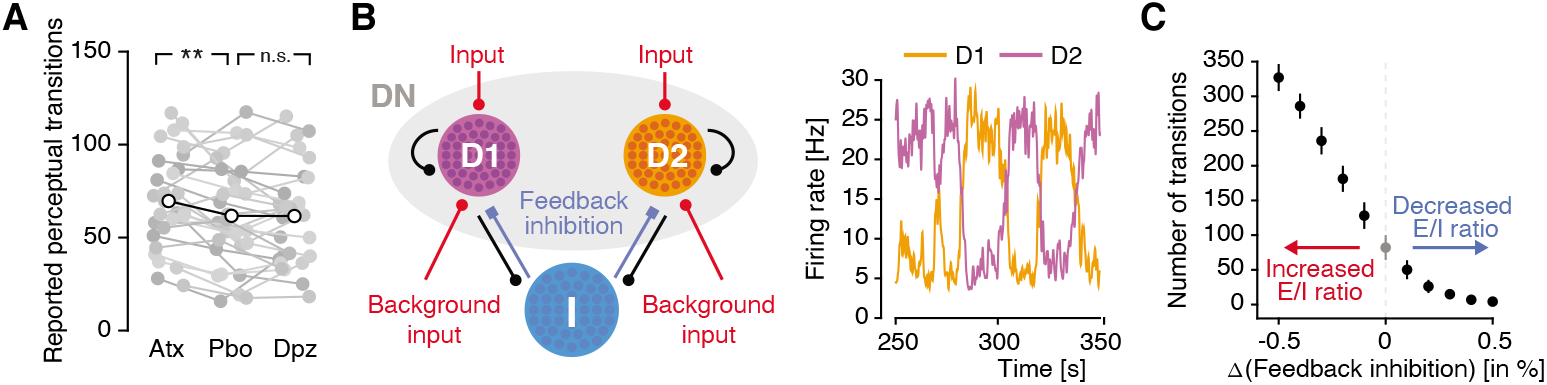
Catecholamine-induced increase in E/I ratio can increase perceptual variability. **(A)** Effect of atomoxetine on rate of alternations in the judgment of continuous input (changes in the apparent direction of rotation of the seemingly rotating sphere). **(B)** Left: Schematic of the decision circuit, endowed with two excitatory decision populations, D1 and D2, and a non-selective population (DN), fully connected to a pool of inhibitory neurons. The two decision populations receive noisy Poisson input, reflecting the ambiguous nature of the visual stimulus. Right: The model exhibits spontaneous firing rate fluctuations. Perceptual transitions in the model are defined as changes in the dominance of one population over the other (i.e., one having a higher firing than the other). **(C)** Effect of E/I increase in circuit model on number of transitions in the judgment of continuous input. E/I increase in the circuit model is implemented via decrease in feedback inhibition (red/blue arrows).

To make the above microcircuit model produce selection behavior, we expanded it by means of two populations of excitatory neurons encoding a specific decision (‘D1’ and ‘D2’), which competed via feedback inhibition (Fig. 3B, left), yielding an architecture equivalent to a well-established model in the context of 2AFC tasks (Wang, 2002). Increasing E/I in this model has been shown to yield more variable decision-making in two-alternative forced choice tasks (Lam et al., 2017). To model our current task, we adjusted some parameters (see Methods) and simulated the model under sustained, and equally strong, input to D1 and D2, modelling unbiased competition. The model exhibited ongoing alternations in the activity dominance of D1 or D2 (Fig. 3B, right) and, just as our participants under atomoxetine, an increase in the alternation rate under increased E/I due to decreased feedback inhibition (Fig. 3C). In other words, participants, as well as the model, were more prone to ‘explore’ different perceptual interpretations of a constant, ambiguous input.

The tradeoff between behavioral exploration and exploitation is commonly studied in other contexts than perceptual multistablity – specifically during foraging for reward in environments with changing reward contingencies (Cohen et al., 2007). An influential view holds that catecholamines render choice behavior more variable in order to facilitate behavioral exploration just when the uncertainty about the environment has increased (Aston-Jones and Cohen, 2005; Cools, 2019; Frank et al., 2009). We, thus, performed a second behavioral experiment to probe the effect of atomoxetine (same dose as for the perceptual task, Fig. 4A) on value-based choice during foraging. We used a modified version of a dynamic foraging task previously used in monkeys (Sugrue et al., 2004) (Methods, Fig. 4B,C). As in the first dataset, atomoxetine increased baseline pupil diameter (Fig. 4D). Participants chose between two visual targets (horizontal/vertical Gabor patches, displayed in different hemifields) which were associated with different reward histories (Fig. 4B,C; Fig. S12B). All but three participants performed the task well, reaching a performance of ~70% (Fig. 4E).

**Fig. 4.**
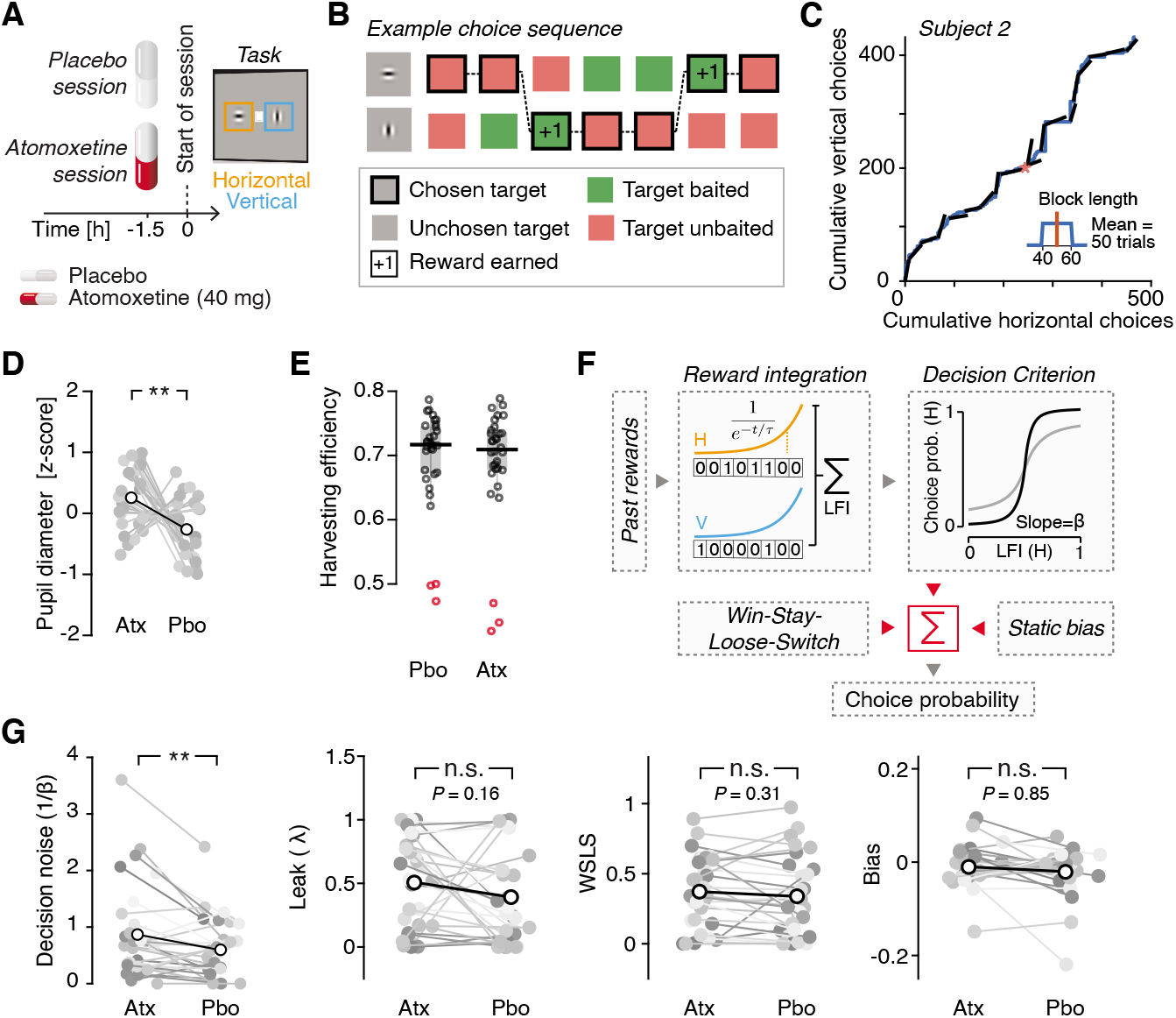
Catecholamines promote exploratory choice during foraging. **(A, B)** Experimental design for value-based choice experiment. **(A)** Administration protocol for the value-based choice experiment: atomoxetine or placebo was administered before each session **(B)** Reward scheme illustrated for example sequence of rewards and choices across seven trials (see Methods for details). **(C)** Choice behavior vs. reward contingencies for an example subject and session. Continuous blue curve, cumulative choices of horizontal vs. vertical targets. Black lines, average ratio of incomes earned from both targets (horizonzal: vertical) within each block). **(D)** Effect of atomoxetine on baseline pupil diameter. **(E)** Harvesting efficiency (fraction of collected over available rewards) per subject and experimental session. Red circles, subjects excluded due to poor performance **(F)** Schematic of the algorithmic model for value-based choice task (dynamic foraging). Choice behavior was analyzed with a reward integration model consisting of four parameters: integrator leak, decision noise (1/β of softmax transformation), weight of ‘win-stay, loose-switch’ heuristic, overall (static) bias (see Methods). **(G)** From left to right: Effect of atomoxetine on the decision noise (1/β), leak (inverse of integration time constant), Win-Stay-Lose-Switch heuristic and bias.

In order to quantify the drug effect on behavioral exploration, we fitted choice fractions with an algorithmic model made up of four parameters that could differ between atomoxetine and placebo (Fig. 4F). The ‘noise’ (1/ß) parameter, governing choice variability (exploration), selectively increased under atomoxetine (Fig. 4G, left). This effect on decision noise was independent of reward integration, which are commonly linked to the dopamine system (Montague et al., 2004). The latter were quantified by the leak (inverse of integration time constant τ, see Fig. 4F), for which we observed no effect (Fig. 4G, second from left). Atomoxetine also did not affect the other two parameters of the model (Fig. 4G). The finding of increased internal noise is in line with the circuit model prediction for E/I increase for forced choice tasks(Lam et al., 2017). In sum, elevated catecholamine levels increased behavioral exploration in sensory- and value-guided behavior, likely via increasing E/I in cortical circuits.

## Discussion

Previous animal work has reported differences between noradrenergic and cholinergic effects on the firing rates or membrane potential fluctuations of single neurons (Castro-Alamancos and Gulati, 2014; Polack et al., 2013). Here, we uncovered a behavioral context-dependent double dissociation between catecholaminergic and cholinergic effects on large-scale cortical network dynamics, developed a mechanistic account of this observation, and validated this account through two independent behavioral tests. Our results may constitute a physiological basis for the distinct roles of catecholamines and acetylcholine in cognition postulated by computational theory (Aston-Jones and Cohen, 2005; Yu and Dayan, 2005). Specifically, a prominent idea holds that the noradrenaline and acetylcholine systems track two forms of uncertainty during inference in changing environments: Acetylcholine signals so-called ‘expected uncertainty’, which originates from the inherent noise corrupting the information received in a given (constant) state of the environment; noradrenaline signals ‘unexpected uncertainty’, stemming from hidden changes in the state of the environment (Yu and Dayan, 2005). These two forms of uncertainty should have separable influences on the neural computations in the cortex that underlie inference. Our results suggest that such functional distinction may be mediated by distinct modulatory effects on local and cortical microcircuits, which in turn translate into massive differences at the level of large-scale cortical interactions.

Our behavioral results establish a general catecholaminergic effect on perceptual and value-based decision-making and confirm a key prediction from a prominent account of catecholaminergic (noradrenergic) modulation of learning and decision-making (Aston-Jones and Cohen, 2005). In this view, the noradrenergic system controls the exploration-exploitation tradeoff, whereby high tonic noradrenaline levels boost behavioral variability. While this is detrimental to performance in static environments, it is adaptive in the presence of hidden environmental changes, as were present in our foraging task, by promoting exploration of alternatives (Aston-Jones and Cohen, 2005; Cohen et al., 2007; Usher et al., 1999). Indeed, animal behavior becomes more variable during periods of high tonic firing of the locus coeruleus in perceptual tasks in static environments (Aston-Jones et al., 1999; Usher et al., 1999) as well in value-based choice in changing environments (Kane et al., 2017; Tervo et al., 2014). In particular, chemogenetic stimulation of locus coeruleus tonic activity, increased decision noise during a foraging task (Kane et al., 2017),, just as in the present Figure 4. These findings in animals are in line with our current results in humans.

One atomoxetine study in humans used a similar pharmacological protocol to ours during a gambling task and decomposed random and directed modes of behavioral exploration (Warren et al., 2017). Unexpectedly, this yielded a *decrease* in random exploration under atomoxetine (Warren et al., 2017), a finding that appears to be at odds with the above animal work as well as the increase in behavioral variability reported in the two tasks of the present study. One possibility is that the predominant effect of atomoxetine on tonic versus phasic noradrenaline level differed between our experiments and the one from (Warren et al., 2017). Such differences may have occurred for several reasons including (i) the different latencies of the behavioral measurements relative to drug intake (1.5 in our study vs. 3 h in theirs), (ii) inter-session differences in baseline arousal/noradrenaline levels, and/or (iii) inter-individual differences in atomoxetine sensitivity between participants. For both our experiments, we found a robust increase of baseline pupil diameter under atomoxetine (Figs. 1B and 4D), consistent with increased tonic noradrenaline levels (Joshi et al., 2016; McGinley et al., 2015; Reimer et al., 2016), an effect not tested by Warren et al. (2017). The decreased exploration in the latter study may have resulted from a predominant increase in phasic noradrenaline release in line with Aston-Jones & Cohen (2005) and also alluded to by the authors (Warren et al., 2017).

In our behavioral model, internal noise (1/β) affected the reward-dependent component of behavior before its combination with WSLS. This was motivated by model comparisons indicating that noise should be applied before, not after, combination of the reward-dependent choice probability with the WSLS heuristic (Fig. S12D). This observation is largely consistent with recent evidence pointing to reward integration, rather than response selection, as the dominant source of behavioral variability (Findling et al., 2019). Separating between noise at each integration step and noise at the transformation from integrated reward (LFI) into choice probability (as in Findling et al., 2019) would require a different modeling approach. Our aim here was to unravel the mechanistic basis of the impact of catecholamines on internal noise, regardless of the exact locus of this noise, based on the monkey work that inspired our task (Corrado et al., 2005; Sugrue et al., 2004). Future work should further constrain the locus of the catecholaminergic noise boost.

Our multi-scale circuit modeling shows that subtle differences in the effects of catecholamines and acetylcholine at the cellular level (e.g. gain increases of different magnitude (Aston-Jones and Cohen, 2005; Herrero et al., 2008; Servan-Schreiber et al., 1990)) can combine to yield context-dependent dissociations at the level of large-scale cortical network dynamics. This principle accounts for the context-dependent (task vs. rest) double dissociation between the modulatory effects observed here. Importantly, our model-based inferences about the underlying circuit mechanisms are consistent with insights from single-cell physiology (Aston-Jones and Cohen, 2005; Froemke, 2015; Haider et al., 2013; Herrero et al., 2008; Martins and Froemke, 2015). The first inference that catecholamines and acetylcholine both increase overall gain (and hence E/I), but with different magnitudes is supported by the observation that noradrenaline and acetylcholine differentially modulate E/I in rodent auditory cortex (Froemke, 2015). Acetylcholine suppresses stimulus-evoked, inhibitory transients in pyramidal cells (Froemke et al., 2007; Letzkus et al., 2011) while noradrenaline suppresses ongoing inhibition in a persistent fashion (Martins and Froemke, 2015). Such synaptic and cellular differences can translate into a differential net gain increase of the whole microcircuit (with a smaller net gain increase under acetylcholine) as described by the nodes of our neural mass model.

Our second inference, of reduced intra-cortical communication (global coupling) under acetylcholine, is also consistent the reduction of intracortical (lateral and/or feedback) signaling that has been observed in in visual and auditory cortex (Hsieh et al., 2000; Roberts et al., 2005; Silver et al., 2008) and in perception (Gratton et al., 2017), possibly mediated by muscarinic receptors(Hsieh et al., 2000). At the computational level, this inference aligns well with the idea that acetylcholine reduces the impact of prior knowledge (intra-cortical signaling) relative to incoming evidence (bottom-up signaling) (Yu and Dayan, 2005). However, evidence from prefrontal cortex suggests that acetylcholine can also increase synaptic efficacy on recurrent intra-cortical connections, through both nicotinic and muscarinic receptors (Arnsten et al., 2010). Further work is needed to elucidate the synaptic basis of the cholinergic effects observed here.

The mechanistic insights put forward here shed new light on apparently inconsistent findings reported in previous studies on pharmacological effects on intrinsic cortical correlations as assessed through neuroimaging (van den Brink et al., 2019). One PET study found cortical an increase in cortical correlations during a task but decrease during rest under clonidine (an a2-adrenergic auto-receptor agonist that reduces noradrenaline release) (Coull et al., 1999), similar to the context-dependence in the present Fig. 1C left. By contrast, a study of atomoxetine (same dose as ours) effects on resting-state fMRI found a robust decrease in correlations (van den Brink et al., 2016), in contrast to weak effect during rest in our present measurements (Fig. 1C, left, lower triangular part). While this may reflect differences in the underlying signals (fMRI vs. band-limited MEG power), our model simulations (Fig. 2) demonstrate how subtle differences in the baseline state of the system (i.e., location on the (*b_E_, b_I_*)-plane) can lead to qualitatively different effects (i.e., sign reversals) of the same gain increase (catecholamines) on intrinsic activity correlations. Differences in environmental factors (e.g. scanner noise), age, or participants’ baseline arousal levels may all shift the baseline state. Such differences in baseline state can translate into qualitative differences of drug effects (including sign reversals) on cortical correlations, even under the same drug and pharmacological protocol (dosage, timing of administration, etc.). This highlights the importance of circuit modeling for understanding the results from pharmacological neuroimaging studies.

Intrinsic correlations in brain activity are widely used in basic human neuroscience and clinical biomarker development (Deco et al., 2014; Fox and Greicius, 2010; Hipp et al., 2012). The behavioral context-dependence of the neuromodulatory effects we uncovered here implies that resting-state measurements alone lack a critical dimension: the comparison between rest and task contexts was necessary to uncover the specific impact of neuromodulators on cortical dynamics.

It is likely that the same holds for other classes of neurotransmitters, and their disturbances in brain disorders. Our approach sets the stage for the development of new non-invasive assessments of the integrity of neuromodulatory systems.

In sum, we have pinpointed candidate circuit mechanisms for the distinct catecholaminergic and cholinergic shaping of large-scale cortical network interactions. Our results can guide future work into the underlying cellular and molecular mechanisms in animals and set the stage for the development of non-invasive biomarkers for the integrity of neuromodulatory systems in humans.

## Acknowledgments

We thank Klaus Wimmer for help with the implementation of the circuit model for decision-making, Christian Büchel, Sander Nieuwenhuis, and Wolf Singer for feedback on an earlier draft of the manuscript, and Alexander Thiele for discussion of cholinergic circuit mechanisms.

## Author contributions

T.P.: Conceptualization, Methodology, Formal Analysis, Investigation, Data Curation, Writing – Original Draft, Writing – Review & Editing, Visualization, Project Administration; A. P.-A.: Methodology, Formal Analysis, Writing – Review & Editing; T.M.: Formal Analysis, Investigation, Writing – Review & Editing; C. G.: Investigation, Writing – Review & Editing; R.L.v.d.B.: Conceptualization, Writing – Review & Editing; G.N.: Provided reagents, Writing – Review & Editing, K.T.: Methodology, Investigation, Writing – Review & Editing; A.K.E.: Resources, Writing – Review & Editing; G.D.: Methodology, Writing – Review & Editing, Supervision; T.H.D.: Conceptualization, Methodology, Writing – Original Draft, Writing – Review & Editing, Supervision;

## Methods

### Pharmacological MEG experiment (resting-state and continuous perceptual choice task)

#### Participants

30 healthy human participants (16 females, age range 20-36, mean 26.7) participated in the study after informed consent. All included participants were non-smokers. The study was approved by the Ethics Committee of the Medical Association Hamburg. Two participants were excluded from analyses, one due to excessive MEG artifacts, the other due to not completing all 3 recording sessions. Thus, we report results from N=28 participants (15 females).

The present dataset was also used in a previous report (Pfeffer et al., 2018), which focused on the effects of both drugs (see below) on the long-range temporal correlations in the local activity fluctuations. The present analyses of the correlations between these fluctuations across different cortical regions are independent from the results presented in this previous work. A different version of the behavioral result shown in Fig. 3A was also shown in the previous paper (Pfeffer et al., 2018).

#### Experimental design

##### General protocol

We manipulated the levels of catecholamines (noradrenaline and dopamine) and acetylcholine through pharmacological intervention (Fig. 1A). Each participant completed three experimental sessions, consisting of drug or placebo intake at two time points, a waiting period of 3 hours, and an MEG recording session. During the recordings, participants were seated on a chair inside a magnetically shielded chamber. Each recording session consisted of six measurement blocks with different behavioral tasks (see below). Each block was 10 minutes long and followed by a short break of variable duration.

##### Pharmacological intervention

We tested for the effects of two different drugs in a doubleblind, randomized, placebo-controlled, and cross-over experimental design. We used the selective noradrenaline transporter inhibitor atomoxetine to boost the levels of catecholamines (noradrenaline and dopamine (Bymaster et al., 2002; Robbins and Arnsten, 2009)). We used the cholinesterase inhibitor donepezil to boost acetylcholine levels. A mannitol-aerosil mixture was administered as placebo. The dosages for both drugs were chosen to be below common clinical steady-state dosages and in accord with previous fMRI work showing clear effects of the same dosages on cortical processing (van den Brink et al., 2016; Silver et al., 2008): 40 mg for atomoxetine (clinical steady-state dose for adults: 80 mg) and 5 mg for donepezil (common clinical entry dose). All substances were encapsulated identically in order to render them visually indistinguishable. Peak plasma concentrations are reached ~3-4 hours after administration for donepezil (Tiseo et al., 1998) and 1-2 hours after administration for atomoxetine (Sauer et al., 2005). In order to maximize plasma drug levels during MEG, participants received two pills in each session, 3 h and 1.5 h before MEG (Fig. 1A): placebo (t = −3 h) followed by atomoxetine (t = −1.5 h) in the ATOMOXETINE condition; donepezil (t = −3 h) followed by placebo (t = −1.5 h) in the DONEPEZIL condition; placebo at both times in the PLACEBO condition. The three sessions were scheduled at least 2 weeks apart to allow plasma levels to return to baseline (plasma half-life of atomoxetine: ~5.2 h – 21.6 h(Sauer et al., 2005); half-life of donepezil: ~70 h).

##### Behavioral tasks

Within each session (and each of the above-defined pharmacological conditions), participants alternated between three different behavioral conditions, all entailing absent or continuous sensory input (2 runs à 10 minutes per condition), here referred to as REST, TASK, and TASK-PRESSING (Fig. 1A, right; see also Supplementary Video 1). REST and TASK were steady-state conditions (absent or minimal variations in sensory input or motor output) tailored to quantifying *intrinsic* correlations between fluctuations in cortical activity. TASKPRESSING was used to validate the behavioral results from the TASK condition. During REST, participants were instructed to fixate a green fixation disk (radius = 0.45° visual angle) in the center of an otherwise gray screen. During TASK and TASK-PRESSING, participants viewed a perceptually ambiguous 3D structure-from-motion stimulus, which was perceived as a rotating sphere (Wallach and O’connell, 1953). The stimulus subtended 21° of visual angle, consisted of 1000 dots (500 black and 500 white, radius: 0.18° of visual angle) arranged on a circular aperture presented on a mean-luminance gray background, and a green fixation dot in the center. In TASK, participants were instructed to count the number of changes in the perceived rotation direction and verbally report the total count at the end of the run. In TASK-PRESSING, the participants were instructed to press (and keep pressed) one of two buttons whenever they perceived a change in the rotation direction. The order of the conditions was as follows for 18 out of 28 participants: (1) REST, (2) TASK-PRESSING, (3) TASK, (4) REST, (5) TASK-PRESSING, (6) TASK. For 10 out of 28 participants the order was reversed: (1) TASK, (2) TASK-PRESSING, (3) REST, (4) TASK, (5) TASK-PRESSING, (6) REST.

The experiment was programmed in MATLAB (The MathWorks, Inc., Natick, United States), using the Psychophysics Toolbox extensions (Brainard, 1997) (PTB-3).

##### Data acquisition

MEG was recorded using a whole-head CTF 275 MEG system (CTF Systems, Inc., Canada) at a sampling rate of 1200 Hz. In addition, eye movements and pupil diameter were recorded with an MEG-compatible EyeLink 1000 Long Range Mount system (SR Research, Osgoode, ON, Canada) and electrocardiogram (ECG) as well as vertical, horizontal and radial EOG was acquired using Ag/AgCl electrodes.

#### Pupil and behavioral data analysis

The pupil diameter recordings were preprocessed as follows: eye blinks as well as eye movements were identified using the manufacturer’s default routines, then padded (+/− 200 ms), linearly interpolated and bandpass-filtered using a second-order Butterworth filter with a passband from 0.01 to 10 Hz. Next, the effect of blinks and saccades on pupil diameter was estimated through deconvolution and removed by means of linear regression (Knapen et al., 2016). Mean pupil diameter was computed in a baseline interval from 6s to 3s prior to the start of each recording block and for all conditions (REST, TASK, TASK-PRESSING). Pupil signals were averaged across the two corresponding blocks. In some cases, pupil diameter was not recorded, or the signal was too noisy. If this was the case for both blocks of a session (placebo, atomoxetine or donepezil), the corresponding subject was not included in the respective analysis. The following number of subjects was excluded/included per combination of conditions: during REST-PLACEBO (N = 0/28), REST-ATOMOXETINE (N=0/28), REST-DONEPEZIL (N = 2/26), during TASK-PLACEBO (N = 0/28), TASK-ATOMOXETINE (N=0/28), TASK-DONEPEZIL (N = 1/27). Behavioral data from TASK and TASK-PRESSING was averaged across the two blocks, resulting in N=28 for all drug conditions for TASK. In the case of TASK-PRESSING, one participant had to be excluded due to missing triggers in the atomoxetine condition, resulting in N=27.

#### MEG signal processing

The MEG signal processing pipeline described is illustrated in Fig. S2A and entailed the following steps.

##### 1) Preprocessing

The sensor-level MEG data were first preprocessed: strong transient muscle artifacts and squid jumps were detected through visual inspection as well as semi-automatic artifact rejection procedures, as implemented in the FieldTrip toolbox (Oostenveld et al., 2011) for MATLAB. To this end, data segments contaminated by such artifacts (+/− 500 ms) were removed from the data (across all channels). Subsequently, the data were downsampled to 400 Hz split into low ([0.5-2]-40 Hz; the lower cutoff was variable across (but identical within) subjects at 0.5, 1 or 2 Hz) and high (>40 Hz) frequency components, using a 4th order Butterworth filter. Both signal components were separately submitted to independent component analysis(Bell and Sejnowski, 1995) using the FastICA algorithm (Hyvarinen, 1999). Artifactual components (eye blinks/movements, muscle artifacts, heartbeat and other extra-cranial artifacts) were identified based on three established criteria (Hipp and Siegel, 2013): power spectrum, fluctuation in signal variance over time (in bins of 1s length), and topography. Artifact components were reconstructed and subtracted from the raw signal and low- and high frequencies were combined into a single data set. On average, 20 (+/− 14) artifact components were identified for the low-frequencies and 13 (+/− 7) artifactual components were identified for the high frequencies.

##### 2) Spectral analysis

From the cleaned MEG signal, spectral estimates were obtained using Morlet’s wavelets (Tallon-Baudry and Bertrand, 1999), similar to previous reports (Hipp et al., 2012; Siems et al., 2016):

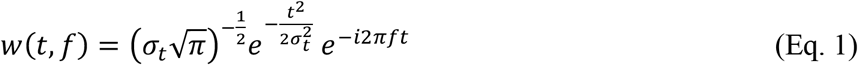

We constructed wavelets for 17 logarithmically spaced (base 2) center frequencies, ranging from 4 Hz to 64 Hz. In keeping with previous work (Hipp et al., 2012; Siems et al., 2016), the spectral band-width was set to half of an octave (*f* /*σ_t_*~5.83) and amplitude as well as phase estimates were obtained for consecutive, half-overlapping segments of a length of ±3*σ_t_*. Segments that contained artifactual samples (see *Preprocessing*) were omitted from the analysis.

##### 3) Source analysis

For the main analyses, we projected the sensor-level signal onto 400 vertices located on the cortical surface, resulting in an estimated source level signal *X_src_*(*r,t,f*). To this end, we estimated source-level power by means of adaptive spatial filtering (linear “beamforming”; Veen et al., 1997), separately for each participant and recording session. For each source location *r* and frequency *f*, a spatial filter *A*(*r,f*) was computed according to:

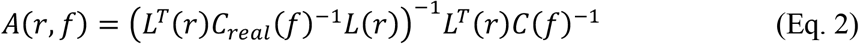

where *L* was the magnetic leadfield, *T* denoted matrix transpose, and *C_real_*(*f*) the real part of the (complex-valued and regularized) cross spectral density (CSD) matrix of the sensor-level data for frequency *f*. *A*(*r,f*) contained three orthogonal projections. We used singular value decomposition of the CSD matrix to determine the direction of the dipole maximizing power (i.e. the first eigenvector) at location *r*. We then computed the corresponding spatial filter for this direction, henceforth referred to as *B*(*r,f*). This filter was used to project the sensor-level data *X*(*t,f*) onto that dominant dipole, as follows:

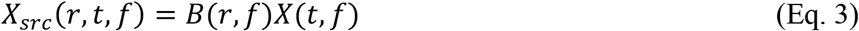

where *X_src_*(*r,t,f*) denoted the complex-valued, source-level spectral estimates for location *r.* Prior to computing the spatial filter, the CSD matrix was regularized with the mean of its diagonal multiplied by a scaling parameter *α*. For the results shown in the main section of this article, this parameter was chosen to be *α* = 0.3 (see Fig. S6 for alternative values of *α*).

##### 4) Orthogonalized power envelope correlations

Inter-regional correlations were computed as the correlations of the power estimates at carrier frequency *f* between two regions *i* and *j,* across all non-artifactual segments. In order to reduce spurious correlations arising from instantaneous signal leakage, we used a procedure established previously (Brookes et al., 2012; Hipp et al., 2012). Specifically, we orthogonalized each signal Y with respect to signal X according to:

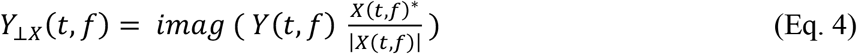

where *YΥχ*(*t,f*) was the signal *Y*(*t,f*) orthogonalized with respect to signal *X*(*t,f*) and * the complex conjugate. Next, the absolute value was taken and the resulting signal was squared, yielding source-level power envelopes, and log-transformed to render the distribution more normal. The orthogonalization was performed in two directions, *Y_Υχ_*(*t,f*) as well as *X_Υγ_*(*t,f*). Correlation coefficients were computed for both directions and the resulting (Fisher transformed) values were averaged. Doing this for all pairs of vertices and for each frequency band resulted in a correlation matrix of size 400×400 for each of the 17 carrier frequencies. In what follows, we refer to these correlation matrices as ‘functional connectivity (FC)’ matrices.

In order to compare the empirical results to results obtained from simulations of a neural mass model (see: *Computational modeling* below), we repeated the above-described procedure for computing source-level FC matrices, but now at coarser granularity. To this end, we selected the 76 cortical regions of the Automatic Anatomical Labeling (AAL) atlas (Tzourio-Mazoyer et al., 2002), excluding the cerebellum and subcortical regions (see Table S1 for included regions). Source locations were first arranged on an equally spaced grid (of 4 mm x 4 mm x 4 mm resolution) covering the entire brain and each grid point was either assigned to one of the 76 selected cortical AAL regions or omitted from further analysis. For each of the vertices that were assigned to one of the 76 AAL regions, frequency-specific source-level estimates for each time point *X_src_*(*r, t,f*) were computed following the procedure outlined above. Next, for each vertex within region *i* (with *i* ∈ {1,…,76}, we computed its average correlation to all vertices of region *j* (after Fisher transformation). This was repeated for all vertices of region *i* after which the correlation values were again averaged across all vertices within region *i.* This procedure was repeated for all 76 regions, resulting in a 76×76 FC matrix for each of the 17 carrier frequencies.

#### Quantification of the topology of MEG correlation structure

##### Degree centrality

We computed frequency-resolved degree centrality *k*(*f*) (i.e., collapsed across all nodes) as well as local degree *k_i_*(*f*) for each of the *i* = 1 … 400 locations (Fig. S2B/C). Degree is defined by the number of edges that connect a given node to all other nodes in the network (Rubinov and Sporns, 2010). To this end, the FC matrices of all subjects (400×400×28) were first submitted to a procedure described previously (van den Brink et al., 2016; Hipp et al., 2012): for each connection between nodes *i* and *j,* where *i* = 1 … 400 and *j* = 1 …400 we assessed if a connection was present as follows: a connection was determined to be present if the correlation between *i* and *j* was significantly larger (*P* < 0.05; two-sided t-test) than the correlations from *i* to all other nodes or from *j* to all other nodes. In case of a present connection, the corresponding entry in the adjacency matrix *A*(*i,j*) was set to 1. If no connection was present, *A*(*i,j*) was set to 0. This was repeated for all possible pairs of vertices and the full adjacency matrix was computed. From this, we computed degree by:

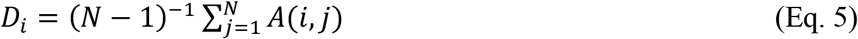

where *N* denoted the number of cortical vertices (N = 400).

#### Statistical tests of MEG effects

##### Cortex-wide changes in cortical correlations

We adopted a previously described two-stage procedure for an unbiased statistical assessment of cortex-wide changes in power envelope correlations (Hawellek et al., 2013). The procedure is illustrated in Fig. S3. The rationale behind the analysis was as follows. Both neuromodulator classes (catecholamines and acetylcholine) might, in principle, increase correlations between some pairs of areas and, at the same time, suppress correlations between other pairs of areas (Eldar et al., 2013). In this case, drug effects might cancel when averaging correlations indiscriminately across all area pairs and comparing average FC between conditions. Instead, our procedure first identified any pairs exhibiting drug-induced increases or decreases above a certain threshold and then tested if the fraction of these pairs was significantly different from what would be expected by chance, separately for pairs with increased and decreased correlations. This procedure was repeated across a wide range of frequencies, yielding the spectra of drug effects shown in Fig. 1E.

For each center frequency *f,* we statistically compared the Fisher-transformed FC matrices, across subjects, between the two drug conditions and placebo, using a two-sided paired t-test. Then we counted the number of significantly positively (*P* < 0.05 and *T* > 0) and the number of significantly negatively altered correlations (*P* < 0.05 and *T* < 0). The resulting value was divided by the number of possible connections M (with M = N*N-N, where N = 400 or N = 76, see above) to obtain the fraction of significantly altered correlations for both effect directions. This procedure was repeated for all 17 frequencies bands (Fig. S3). We employed a single threshold permutation procedure to derive P-values that accounted for multiple comparisons across frequencies (Nichols and Holmes, 2002). For each of *N_p_* = 10000 permutations, the experimental labels (drug conditions) were randomly re-assigned within subjects and the aforementioned procedure was repeated. This resulted in a *N_p_* x17 matrix for both effect directions (significantly increased and significantly decreased correlations). Next, for each permutation, the maximum value across all frequencies (independently for increased and decreased correlations) was determined, yielding a maximum permutation distribution. In order to derive P-values, the empirical results were now compared to this maximum permutation distribution. This procedure is analogous to a single-threshold permutation test commonly applied in fMRI (Nichols and Holmes, 2002), with the single-threshold test being performed across frequencies instead of space (i.e., voxels). In order to test the robustness of the obtained results, we repeated the procedure described above using various alpha values for the initial paired t-test, ranging from a=0.01 to a=0.10, which led to numerically different, but qualitatively similar results (Fig. S7). Through applying an initial thresholding (t-test), spurious and weak changes in correlations are less likely to contribute to the observed result. The initial t-test thus ensures that only changes that are somewhat robust are taken into account.

Significant alterations in the correlations between two regions can be achieved in different ways. A decrease in correlations, for instance, can mean that a positive correlation becomes weaker, or that a negative correlation becomes more negative. However, only the former would qualify as a meaningful reduction in correlation, whereas the latter correlation gets numerically smaller (i.e., more negative), but stronger in terms of the linear dependence between two signals. Hence, a “significant decrease” does not always carry the same meaning, and the same is true for increases. In this data set, the number of positive correlations by far outnumbered the number of negative correlations. In fact, in the alpha and beta frequency range, where the main effects for atomoxetine and donepezil are observed, more than 90% of all connections were positive (across all blocks, and contexts; placebo only). Thus, we interpret an increase (decrease) in correlation in terms of a positive correlation becoming stronger (weaker).

##### Cortex-wide changes in local cortical variability

Changes in the correlation between two signals can be driven by changes in their covariance (numerator of correlation coefficient) as well as changes in the variance of one or both of the signals (denominator). In order to rule out this possibility, we tested for drug-related changes in the variance of local power envelopes across frequencies (Fig. S5). To this end, we have adopted a procedure similar to the one employed to assess changes in cortex-wide activity correlations. First, we computed the variance of the power estimates across half-overlapping temporal segments (see *Spectral estimation),* separately for each of the 17 carrier frequencies. Next, we counted the fraction of nodes that exhibited significantly altered variance, separately for increases and decreases. We employed the same permutation procedure described above in order to derive corresponding permutation distributions from which P-values were computed (two-sided single threshold permutation test). Analogous to the ‘fraction of significantly altered correlations’, this procedure yielded, per frequency band, the fraction of vertices (nodes) with significantly positively or negatively altered variance.

#### Cortical circuit modeling

##### Large-scale neural mass model

###### Single node dynamics

We simulated neural population activity using a mean field model based on the Wilson-Cowan (WC) equations (Deco et al., 2009; Wilson and Cowan, 1972). Each local WC node consists of an excitatory and an inhibitory neuronal population (Fig. 2A). The dynamics of the E and I populations of each node are governed by the following stochastic differential equations:

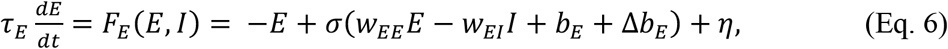

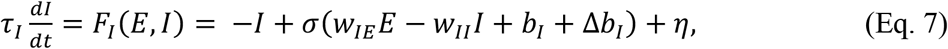

where *E* and *I* represent the firing rates of excitatory and inhibitory populations, respectively. Since we were interested in neural oscillations, the model parameters were chosen to generate oscillatory dynamics. The local synaptic weights interconnecting the excitatory and inhibitory populations were given as *w_EE_* = 12, *w_IE_* = 16, *w_II_* = 4 and *w_EI_* = 12. *b_E_* and *b_I_* represented external background inputs to the he excitatory and an inhibitory, respectively, Δ*b_E,I_* represents task-induced input (Δ*b_E_* = Δ*b_E_* = Δ*b_I_* 0, for resting dynamics), and *η* was uncorrelated Gaussian noise with amplitude equal to 0.005. Time constants were set to *τ_E_* = 9 ms and *τ_I_* = 18 ms for excitatory and inhibitory populations, respectively. The (non-linear) transfer function converting input currents into output firing rates, *σ*(*u*), was chosen to be a sigmoid:

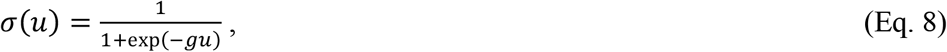

where *g* determined the slope of the input-output function for both excitatory and inhibitory populations (i.e., response gain of the node).

The solutions (*E*\*,*I*\*), or fixed points, of the coupled equations 6 and 7, were given by *E*\* = *σ*(*w_EE_E*\* – *w_EI_I*\* + *b_E_*) and *I*\* = *σ*(*w_IE_E*\* – *w_II_I*\* + *b_I_*), yielding solutions depending on the external inputs (*b_E_,b_I_*), which are the bifurcation parameters of the system. In the (*b_E_, b_II_*) parameter space, we observed a region of noise driven oscillations (i.e., a spiral; damped oscillations that, in the presence of noise, result in noisy oscillations) and a region of sustained oscillations (i.e., a limit cycle) (Fig. S8A).

###### Two coupled nodes

We first studied the effect of gain modulation on correlations during REST and TASK in a minimal network composed of two WC nodes (Fig. 2B, left). This step will provide intuitions before studying the whole-brain network composed of 76 nodes interconnected through a connectome. Let the excitatory populations of the nodes be connected through a reciprocal coupling *c* (in the two-node model, *c* = 1; see below for cortex-wide model). The firing rates of node 1 evolve as:

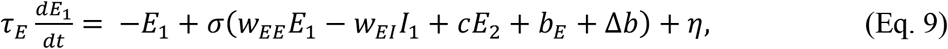

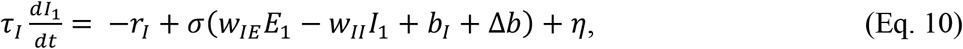

and analogously for the firing rates, *E*_2_ and *I*_2_, of node 2.

To study the correlations between nodes in the parameter space, we used a linear noise approximation described in detail in the Supplementary Information. Using this approximation, we studied how changes in gain, i.e., *g* → *g* + Δ_*g*_, and inputs, i.e., (*b_E_, b_I_*) → (*b_E_* + Δ*b, b_I_*, + Δ_*b*_), change the correlation between the two excitatory populations both during REST (Δ*b =* 0) and TASK (Δ*b* ≠ 0). In this way, we can test hypotheses on the parameter changes induced by ATOMOXETINE and DONEPEZIL, assuming that TASK changed the background inputs *b_E_* and *b_I_*, and the drugs changed *g*. Note that in the case of the two-node model the task-related change of the background inputs was equal for E and I, i.e., Δ*b_E_* = Δ*b_I_*, = Δ*b*.

In sum, the change in correlation between excitatory populations during REST was given as:

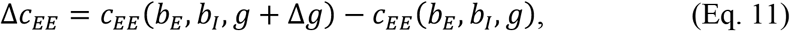

and the change in correlation between excitatory populations during TASK was given as:

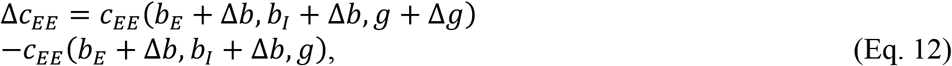

Fig. 2C maps the change of correlations during REST under gain modulation in the (*b_E_, b_I_*)-plane. The combined effect of drugs on parameters *b_E_* and *b_I_*, and the effect of TASK, were obtained by translating the state of the system in this map.

###### Cortex-wide model

In order to directly compare the computational model to the empirical results, we simulated a cortex-wide variant of the model. For each of 76 cortical AAL nodes, the dynamics were governed by the following differential equations:

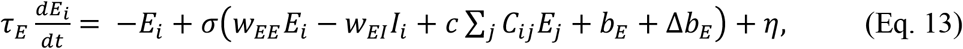

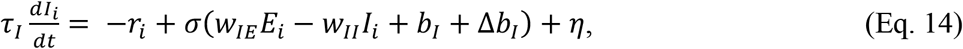

where *i,j ϵ* {1,2,…,76}. In this cortex-wide model, an additional parameter was incorporated: long-range cortical connectivity between all possible pairs of regions, given by *C_ij_*, which is scaled by the global coupling parameter *c*. The matrix *C_ij_* was given by a structural connectivity matrix used in previous studies (Deco et al., 2017, 2018) and was estimated by means of diffusion tensor imaging (DTI). Details can be found in the respective publications.

For the simulation, equations 13 and 14 were integrated using the Euler method with dt = 0.01.

The model was run for a wide range of background inputs to excitatory (*b_E_*) and inhibitory populations (*b_I_*). In order to assess activity correlations (functional connectivity) in the model, we computed all-to-all (76×76) pairwise correlations between the raw time series of excitatory firing rates *E_i_*. The model was simulated, for a total of 58.5s for each parameter combination prior to each run, initial conditions were randomized and the first 1.8 seconds were excluded from further analysis in order to avoid transient effects due to the initial conditions. All simulations and corresponding analyses were carried out in MATLAB2019b.

###### Identification of the oscillatory regime in the cortex-wide model

In all model-based analyses, we assume that the (healthy) human brain never resides in a dynamical regime of sustained oscillations (Fig. S8A, top panel; see also Supplementary Information). We therefore identified, and excluded, parameter combinations resulting in sustained oscillations in the cortex-wide model (see above). In order to identify parameter combinations where simulated population activity settled in the oscillatory regime, the cortex-wide model was simulated *without noise* (i.e., *η* = 0) for a total of 58.5s (plus 1.8s initialization, as outlined above). Next, for non-overlapping segments of 27 ms, the maximum and minimum of *r_E_* was computed. In a regime of noise-driven (damped) oscillations, the activity relaxed back to a fixed point over time (Fig. S8A, middle and bottom). Hence, the computed maximum and the minimum should converge on the same value, whereas in a regime of sustained oscillations, the maximum and the minimum will remain different throughout the entire simulation (Fig. S8A, top). Consequently, the regime was defined as non-oscillatory or *noise driven*, if: (1) the maximum and minimum were identical at any point in time or (2) the difference between maximum and minimum decreased monotonously over time (indicative of a damped oscillation); if none of the two were true, the signal was defined as a sustained oscillation (see Supplementary Information).

###### Model fitting procedure

We fit the free parameters of the cortex-wide model through an iterative procedure. The purpose of this procedure was to identify two working points, mimicking the two behavioral conditions (REST and TASK). First, we estimated the global coupling parameter *α*. This this end, we simulated the cortex-wide model (76 regions) over a range of 41 different coupling parameters *α* (with *α ϵ* {0,0.05,…,2}) and across 61×61 combinations of background inputs (with *I_E_ ϵ* {−4, −3.9, …, −1} and *I_I_ ϵ* {…5, −4.9, …, −2}). We then estimated the similarity of the simulated functional connectivity matrix *FC_sim_* and the empirical functional connectivity matrix *FC_emp_* (Rest and Placebo only; averaged across frequencies that showed significant changes for both drugs; see Fig 1E), separately for each combination of *I_E_* and *I_I_*, by means of a distance metric *δ* based on Pearson correlation(Demirtaş et al., 2019a):

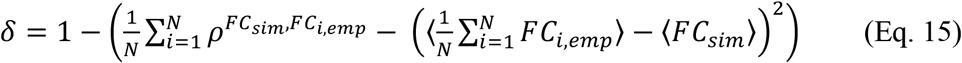

where *ρ^FC_sim_,FC_i,emp_^* was the correlation (i.e., pattern similarity) between the empirical FC matrix for subject *i* (averaged across frequencies, with *i ϵ* {1,2,…,28}) and the simulated FC matrix and 〈 〉 denotes the average across all possible connections. We averaged the resulting distance values *δ* across all external background inputs (*b_E_* and *b_I_*), while omitting parameter combinations where the network activity settled into a regime of sustained oscillations (see above). This resulted in a mean distance 〈*δ*〉 for each level of global coupling *α* (Fig. S8C). Additionally, we repeated the procedure but instead computed Pearson correlation between *FC_sim_* and *FC_emp_* (Fig. S8D). We identified the level of *α* where the mean distance 〈*δ*〉 between *FC_sim_* and *FC_emp_* was minimized (*α* = 1.2). The parameter with lowest 〈*δ*〉 also yields a high correlation between *FC_sim_* and *FC_emp_* (Fig. S8D).

After having fixed the global coupling parameter, we aimed to identify the combination of *b_E_* and *b_I_*, for each individual participant, that resulted in the highest similarity between *FC_sim_* and *FC_emp_* during REST and PLACEBO (i.e., lowest distance). To this end, we first identified the combinations of *b_E_* and *b,* where the distance between *FC_sim_* and *FC_emp_* was below the 2.5^th^ percentile. The resulting binary matrix was then submitted to a clustering procedure (using SPM’s ‘bwlabel’ function) and the single largest cluster was extracted. This was to reduce the influence of spurious correlations on the fitting procedure. Next, the geometric center of the largest cluster was computed and defined as the best fitting combination of *I_E_* and *I_I_*, yielding a working point for REST_sim_. This procedure was repeated separately for each of the 28 participants (Fig. S8F). In order to determine the corresponding TASK parameter TASK_sim_, we assumed that the constant visual stimulation during TASK increases both excitatory (*b_E_*) and inhibitory drive (*b_I_*,), consistent with electrophysiological recordings in rodent visual cortex V1(Adesnik, 2017; Haider et al., 2013) (see Supplementary Information). Thus, in order to simulate TASK (TASK_sim_) we increased the background input to both excitatory and inhibitory populations, i.e., *b_E_* and *b_I_*. We chose to increase background input to inhibitory populations by Δ*b, =* 0.475 and to excitatory population by Δ*b_E_* = 0.25 (Fig. 2D,E, cortex-wide model). Note that, for simplicity, we here assume that the change in background inputs due to TASK is global and homogenous across all nodes and identical for all participants. This assumption is certainly oversimplified and model fits can likely be improved by heterogeneous scaling of these effects as well as by taking individual differences into account.

##### Local microcircuit models

###### Microcircuit model of local node

In order to assess how changes in neural gain can be achieved through specific changes in synaptic weights, we simulated a model of a canonical cortical microcircuit, as a conductance-based neural network (Fig. 2F) comprised of 400 leaky integrate-and-fire units (20% inhibitory). Model equations and parameters follow (Wang, 2002), with some modifications as mentioned below. The model architecture is depicted in Fig. 2F. The membrane potential dynamics of the excitatory units below threshold were governed by:

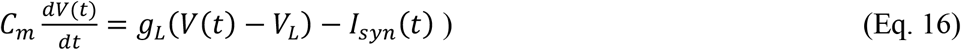

Here, *I_syn_*(*f*) denotes the total synaptic current, which was composed of two glutamatergic excitatory currents (with AMPA and NMDA components) and GABAergic inhibitory currents. External input as well as external noise to the network were mediated exclusively via AMPA receptors. Baseline parameters were identical to the original version (Wang, 2002), with the exception of *g_IE,GABA_* = 1.99 (weight of inhibitory to excitatory synapses) and *g_ext,AMPA_* = 2.5 (weight of external input on excitatory neurons). Moreover, the rate of the external Poisson input to excitatory and inhibitory neurons was changed to *v_ext_* = 881 *Hz* (originally *v_ext_* = 2400 *Hz*). From these baseline values, we parametrically scaled the conductance parameters *g_IE,AMPA_* and *g_EE,GABA_* (AMPA-mediated recurrent excitation and feedback inhibition, respectively) in order to achieve plausible spontaneous dynamics (see Supplemental Information and Fig. S10 for details). Next, we presented the network with stimuli in form of external excitatory input (added to the background input) to all excitatory cells, mediated through AMPA receptors, and assessed the effect on resulting excitatory population firing rate (Murphy and Miller, 2003). In visual cortex, neurons respond to stimuli with increasing contrast with higher firing rates. This relation between a neurons output and the visual input strength is well-described by a hyperbolic ratio function known as the Naka-Rushton function:

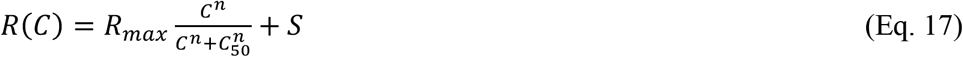

where *R*(*C*) is the firing rate at input contrast *C, R_max_* is the response gain, .*S* reflects the level of the spontaneous (background) activity and *C*_50_ is the stimulus strength that yields a firing rate at half the maximum. Using this equation, we generated a set of stimuli (with varying “contrast”, i.e., varying levels of *C*) that were transformed into firing rates of different frequency and were subsequently fed into the network as an AMPA-mediated excitatory Poisson input. The parameters used in the current study were identical to the parameters used in a previous theoretical study on the effects of excitation and inhibition on response gain of single neurons (Murphy and Miller, 2003): *R_max_* = 2000 *Hz*, *C* = 20.133, *n* = 1.2 and .*S* = 0. This approach allowed us to measure the response of a neural population to inputs of varying contrast strength, which is typically depicted as a contrast-response curve (Fig. 2F, bottom panel). In order to assess the effect of excitation-inhibition ratio on the shape of the contrast-response curve, we either decreased or increased feedback inhibition in the model, through adjusting *g_EI,GABA_* (see Supplemental Information for details). Using nonlinear least squares estimation, we fit the hyperbolic ratio function (Eq. 17), with four free parameters (*R_max_*, 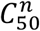, *S* and *n*), to the resulting contrast-response curves. This yielded, among others, response gain parameters (*R_max_*) for different levels of feedback inhibition (Fig. 2F, bottom panel). The network was simulated for 3s per parameter combination, with a similar period of external stimulation. All simulations and analyses were carried out in Python 2.7.15, using the Brian spiking neural network simulator (version 1.4.4) (Goodman, 2009; Goodman et al., 2014), the Elephant toolbox for Python as well as custom code. The Python code for the model simulations was adapted from publicly available code (Wimmer et al., 2015).

###### Decision circuit

In order to understand how the increase in reported perceptual transitions during ambiguous visual stimulation under atomoxetine (Fig. 3A; Fig. S11) could be related to changes in synaptic activity, we extended the above neural circuit by equipping the model with two excitatory populations, which competed for dominance via common feedback inhibition. The synaptic equations were identical to the homogeneous microcircuit described in the previous section. Unless stated otherwise, model architecture and parameters were identical to the original description (Wang, 2002) (Fig. 3B, left). The circuit consisted of N=2000 leaky integrate-and-fire neurons, endowed with full connectivity. 1600 of the neurons were excitatory and 400 inhibitory. The excitatory cells were assigned to one of three subpopulations: two decision populations (240 neurons each), D1 and D2, as well as one non-specific population (DN; 1120 neurons). The two decision populations were assumed to represent the populations that encode the two possible perceived rotation directions of the ambiguous stimulus. All neurons, excitatory and inhibitory, of all populations (D1, D2, DN and I) received independent AMPA-mediated excitatory background input in the form of a Poisson spike train with a frequency of 2880 Hz. In addition, the neurons of the decision populations D1 and D2 received independent AMPA-mediated excitatory input with a mean firing rate of 55.6 Hz which was to reflect the stimulus-related sensory input. The identical mean in input to both decision populations was to mimic the ambiguous nature of the structure-from-motion stimulus. Recurrent connections within D1 and D2 were stronger than connections within DN, by a factor of w_+_ = 1.6. The network was simulated for 600s and population firing rates were estimated for time bins of 100 ms length. Perceptual transitions in the model were defined as the time points where the firing rate of one decision population exceeded the firing rate of the other decision population, i.e., at those time points where the difference between firing rates of D1 and D2 changed in sign (Fig. 3B, right). In order to attenuate the effect of very fast fluctuations on the number of perceptual transitions, we low-pass filtered the firing rates of both decision populations prior to computing the perceptual transitions (cutoff frequency 1 Hz). In order to understand the effect of execution and inhibition on perceptual transitions, we again modified feedback inhibition by means of adjusting *g_EI,GABA_* and computing the number of perceptual transitions for each level of feedback inhibition. For each level of feedback inhibition, the network was simulated 20 times.

#### Pharmacological behavioral experiment (value-based choice task)

##### Participants

We measured 32 participants (21 females, age range 20 – 36, mean 27.28) that performed two sessions of a value-based choice task (Fig. 4A; Fig. S12A) after informed consent. All included participants were non-smokers. The study was approved by the Ethical Committee responsible for the University Medical Center Hamburg-Eppendorf. We excluded three participants from the analysis based on foraging efficiency, which we here defined as the fraction of collected rewards over the total number of available rewards: we excluded participants whose foraging efficiency deviated more than three times the median from the median, scaled by a constant (c≈1.4826, using MATLAB’s ‘isoutlier’ function). Based on this criterion, the same three participants were excluded for both experimental sessions (Fig. 4E). This resulted in in 29 included participants.

##### Experimental design

###### General protocol

We manipulated the levels of catecholamines (noradrenaline and dopamine) in a double-blind, randomized, placebo-controlled pharmacological intervention using atomoxetine (see above, section *Pharmacological MEG experiment).* Each participant completed two experimental sessions, consisting of drug or placebo intake, a waiting period of 1.5 h, and performance of the behavioral task during MEG recordings. During task performance, participants were seated on a chair inside a magnetically shielded chamber and the (visual) task stimuli were presented on a screen in front of them (Fig. 4A; Fig. S12A). Because this was a standard trial-based task design entailing many sensory and motor transients, the MEG data from this task were not used for the analysis of correlations between intrinsic fluctuations in cortical activity. The MEG data will be reported in a separate study.

###### Behavioral task

We used a modified version of a dynamic foraging used in a previous monkey physiology study (Sugrue et al., 2004). Participants chose freely between two visual target stimuli (identified by orientation, randomized by position), which were associated with different histories of monetary rewards. The sequence of events during each trial is shown in Fig. S12A. Participants were asked to fixate a white box in the center of a uniform grey background. Each trial started with the presentation of the two targets (full-contrast Gabor patches with vertical or horizontal orientations) that were presented on either side of the fixation mark (eccentricity ~8.5°, diameter ~4.25° visual angle). The horizontal target’s (left vs. right) location was randomly drawn on each trial, under the constraint that it would appear equally frequently on each side within a block of trials with equal ‘income ratio’ (see below). After a 0. 5-1.5 s delay, the fixation mark changed shape (from box to diamond), prompting the subject’s choice. Participants then pressed a button with their left or right index finger to choose the target at the corresponding location. After another variable delay (2-5 s), subjects received auditory feedback on the outcome of their choice (reward or no reward) by means of a low- or high-pitched tone (low: 200 Hz, high: 880 Hz; each with duration: 150 ms). The mapping of the tones to ‘reward’ or ‘no-reward’ was counterbalanced across participants and instructed before the start of each experimental session. If the participant had not yet responded within a deadline of 3 s, another tone (440 Hz, 50 ms) signaled their missed response (no reward), and the trial was aborted. Targets disappeared after feedback tone, so that only the fixation mark remained for an inter-trial-interval (ITIS) 2-5 s, during which the subjects kept fixating. The next trial started upon the new onset of the targets. Trial duration varied between 4.5 and 11.5 s plus reaction time (reaction times could range between 0 and 3 s) respectively, with an average trial duration of 8 s plus reaction time (0 to 3s). Participants completed 525 trials in each experimental session, taking about 75 min, excluding breaks in between blocks of 100 trials.

Each target was baited with a separate Poisson process for generating rewards, under the following constraints (Fig. 4B,C): (i) the ‘income’ (i.e., reward) rate averaged across both targets was 0.8 rewards per trial; (ii) the ratios between the reward rates associated with each target for a given block of trials (see below) were drawn from a predefined set {7:1, 5:1, 3:1, 1:1, 1:1, 1:3, 1:5, 1:7}; (iii) a reward assigned to a target (i.e., orientation) remained available there until this target was chosen; (iv) when a reward was available at a target, no new reward could become available there (i.e., there was never more than one reward available per target). Correspondingly, both or one or none of the targets could carry a reward in a given trial – the rewards associated with both targets were uncoupled.

The ratios between reward rates (‘local income ratios’; Sugrue et al., 2004) changed between blocks of trials, without this being signaled to the participants. The block duration was sampled from a uniform distribution which ranged between 40 and 60 trials (Fig. 4C). Subjects were not informed about these changes. Because of this dynamic nature of the foraging task, a successful policy is to integrate rewards earned from choosing each target, but only ‘locally’ in time, over the last trials (see Sugrue et al., 2004, and *Behavioral modeling*)

Subjects were not instructed about the statistics of the process generating the rewards. They were only instructed to (i) try to earn as many rewards as possible and that this would translate to a bonus payment at the end of the session; and (ii) to be ‘flexible’ in their behavior because the relative income of the two targets could change over time.

Subjects were rewarded €0-20 bonus based on performance. The lower boundary was chance level performance; the maximum bonus could be earned by performing on par with an ideal observer model, which chose based on full information about the reward ratio at every trial.

##### Pupil analysis

The pupil diameter recordings were preprocessed similar to experiment 1 (see *Pharmacological MEG experiment):* eye blinks as well as eye movements were identified using the manufacturer’s default routines, then padded (+/− 200 ms), linearly interpolated and bandpass-filtered using a second-order Butterworth filter with a passband from 0.01 to 10 Hz. Next, the effect of blinks and saccades on pupil diameter was estimated through deconvolution and removed by means of linear regression (Knapen et al., 2016). Mean pupil diameter was computed in a pre-target baseline interval from 500 ms to 0 ms prior to target onset. Pupil recordings were not available for 4 participants. Hence, the analysis was performed for the remaining 25.

##### Behavioral modelling

We fitted an algorithmic model of behavior to quantify the effects of atomoxetine on the different computations governing decision-making in the task. Our model extended the model previously developed to account for monkey choices in the task(Corrado et al., 2005; Sugrue et al., 2004). A schematic of the model is depicted in Fig. 4E. In words, model choices were computed through the following steps: (i) leaky integration of the rewards gathered from choosing each option over the recent trials (locally in time); (ii) combination of the ‘incomes’ earned from each reward into a relative value signal, the ‘local factional income’ (LFI); (iii) non-linear (*softmax*) transformation of LFI into a probability of choosing the horizontal option; (iv) a weighted contribution of a winstay-loose-switch (WSLS) heuristic and (v) a weighted contribution of general bias (preference for one of the targets) to the final choice probability. Leaky reward integration was applied in order to account for rapid adaption to the hidden changes in income ratio across blocks (see above: *Experimental design* and (Corrado et al., 2005; Sugrue et al., 2004)).

Please note that the WSLS heuristic had been suppressed, by design, in Sugrue et al. (2004)(Sugrue et al., 2004) through a so-called ‘change-over-delay’ (i.e., punishment for switching targets after one choice). We did not include this change-over-delay in our task to render the foraging task even more naturalistic. We found that subjects’ behavior could be well accounted for by a linear mixture of the leaky reward integration described by steps (i)-(iii), the heuristic from step (iv) (Fig. 4F).

In line with Sugrue et al., 2004, we fitted the model by minimizing the negative log-likelihood between the model choice probability from step (iv) and the subjects’ binary choices, giving the set of parameter values (for similar approach see Sugrue et al., 2004). We first found the minimum in a rough grid search. These parameter values were then used as starting point for MATLAB’s ‘fminsearchbnd’.

For each trial *t*, the model computed *LI_hor_*, the ‘local income’ earned from choosing the horizontal option, as follows:

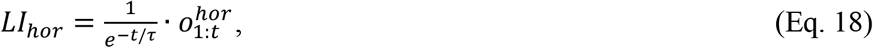

where 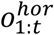 were the outcomes of horizontal choices on trials 1: *t* and *τ* was the reward integration time constant (model leak 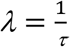). Rewards earned from choosing the horizontal option were coded as 1 and all non-rewarded horizontal choices, or choices to the vertical option (irrespective of reward) were coded as 0. The same equation was used to update *LI_ver_*, now coding rewards earned from choosing the vertical option as 1 and all other outcomes as 0.

The local fractional income *LFI_hor_* was defined as:

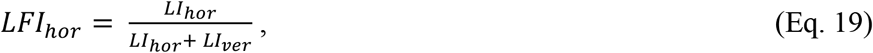

*LFI_hor_*was transformed into the choice probability *p*(*c_t_* = *hor*), defined with respect to the horizontal target, through a sigmoidal (softmax) function (Corrado et al., 2005):

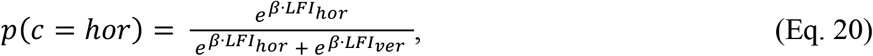

where *β* was the inverse temperature parameter that governed decision noise, i.e. corrupting the mapping from LFI to the behavioral choice. *β* ranged from 0 to infinity (no noise). This placement of the softmax transformation was motivated by model comparison (Fig. S12D).

Choice probability *p*(*c_t_* = *hor*) was further transformed by linear combination with the simple switching (WSLS) mechanism defined as follows:

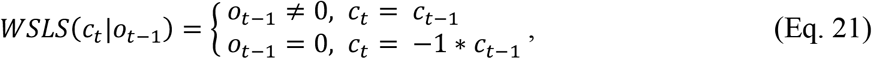

Finally, *p*(*c_t_* = *hor*) was transformed into the final choice probability estimate by linear combination with general bias *δ*, which could range between −1 (all choices to vertical option) to 1 (all choices to horizontal option).

In sum, the dynamics of choice probability (model quantity fitted to the data), was given by:

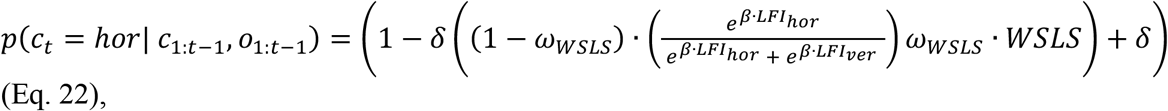

where *p*(*c_t_* = hor | *c*_1:*t*−1_, *o*_1:*t*−1_) was the probability of horizontal choice on trial *t*, given the choices made and outcomes (rewards) received from trial 1 to trial *t*-1, *ω_WSLS_* was a free parameter (ranging from 0 to 1) that controlled the contribution of the WSLS heuristic to choice probability.

In the results reported here, the model included all four free parameters for each of the participants. The level of bias and tendency to rely on the WSLS heuristic varied substantially between participants. In a separate version of the analysis, where we determined the best-fitting set of parameters per participant using cross-validation, we replicated the increase of the softmax parameter under influence of atomoxetine (data not shown).

## Supplementary Information

### Supplementary Methods

#### Dynamics of a single-node Wilson Cowan model

The mass dynamics of the local nodes were governed by the control parameters and ***b_E_*** and ***b_I_***, representing external drive (background input) to the excitatory and inhibitory populations, respectively. The parameters of the model were tuned such that the model is in a dynamical state close to a so-called supercritical Adronov-Hopf bifurcation. The ‘Hopf-bifurcation’ separates a regime in which the system relaxes to a stable fixed point, or *focus*, by drawing a spiral in the phase space (noise-driven, damped oscillations; Fig. 2C; Fig. S8A, middle and bottom), and a regime in which the network activity settles into a *limit-cycle,* a closed orbit in the phase space corresponding to a periodic solution (sustained oscillations; Fig. 2C and Fig. S8A, top). When the system settles into the focus, intrinsic noise induces stochastic oscillations and gives rise to a broad spectral density with a single peak. In contrast, in the limit-cycle, autonomous regular oscillations are observed, with a spectral density presenting a narrow peak (Fig. S8E). The parameters *w* as well as ***b_E_*** and ***b_I_***, were loosely adjusted in order to produce dynamics in the vicinity of a supercritical Hopf bifurcation. In the context of the Hopf-bifurcation, the term ‘supercritical’ is not to be confused with the same term referring to self-organized criticality and power law scaling behavior (Beggs and Plenz, 2003; Poil et al., 2012).

#### Linear noise approximation for two-nodes Wilson Cowan model

We used a linear noise approximation to study the linear fluctuations around the system’s fixed points, i.e., 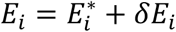 and 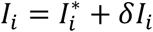, where the fixed points are given by:

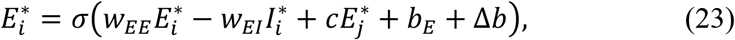

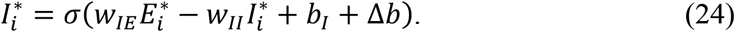

Dynamic equations for the linear fluctuations can be written as:

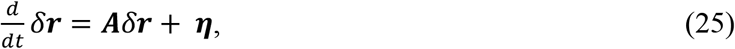

where *δ**r*** = [*δE*_1_, *δI*_1_, *δE*_2_, *δI*_2_], ***η*** is the noise matrix, and ***A*** is the Jacobian matrix of the system evaluated at the fixed points, given by the 4-by-4 matrix:

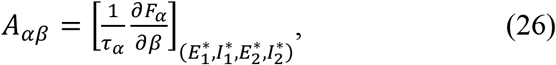

where *α,β* ∈ {*E*_1_, *I*_1_, *E*_2_, *I*_2_}. Noting that *σ*′(*u*) = *gσ*(*μ*)[1 – *σ*(*u*)] and that, by symmetry, 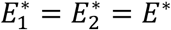 and 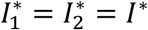, we get:

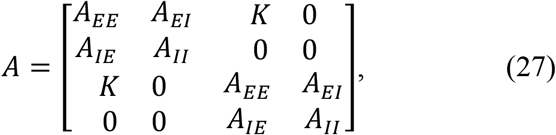

where

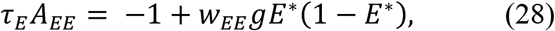

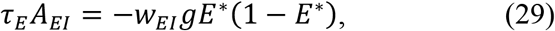

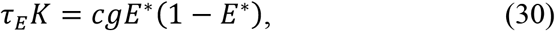

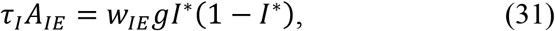

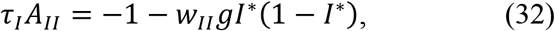

The stationary covariances ***C***_*v*_ = 〈*δ**r**δ**r**^**T**^*〉 between all populations can be obtained through the Jacobian matrix, by solving the following equation:

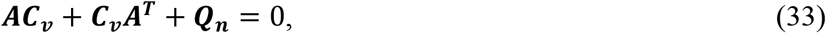

where ***Q_n_*** = 〈***ηη***^***τ***^〉 is the covariance matrix of the noise (which is diagonal for uncorrelated white noise) and the superscript *T* denotes the transpose operator. Note that the Jacobian matrix depends on the system’s fixed points, i.e., it depends on the state of the nonlinear system and, thus, on the external inputs (*b_E_,b_I_*). Hence, the correlations are also a function of the parameters (*b_E_,b_I_*). Equation 20 can be solved using the eigen-decomposition of the Jacobian matrix evaluated at the fixed points: *A* = *LDL*^−1^, where *D* is a diagonal matrix containing the eigenvalues of *A,* denoted *λ_i_*, and the columns of matrix *L* are the eigenvectors of *A.* Multiplying Equation 20 by *L*^−1^ from the left and by *L*^−†^ from the right (the superscript dagger being the conjugate transpose) we get:

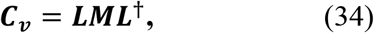

where ***M*** is given by: 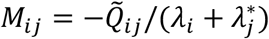 and 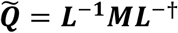.

### Supplementary Discussion

#### Assumptions for large-scale (Wilson Cowan) modeling

Our large-scale modeling approach was based on two assumptions. In the following, we discuss the physiological evidence supporting these assumptions.

##### Assumption 1. Cortex operates in a regime of noise driven, not sustained, oscillations

We assumed that the cerebral cortex generally operates in a regime of noise-driven oscillations, rather than self-sustained oscillations (Fig. 2C; Fig. S8A). In the noise-driven (also referred to as fluctuation-driven) regime, stochastic fluctuations in activity drive damped oscillations in the local nodes. Superposition of such damped oscillations, triggered at random moments in time, give rise to the same ongoing variations in the amplitude of band-limited activity that are commonly observed in electrophysiological data under steady-state conditions (Hipp et al., 2012; Leopold et al., 2003), including the current data set (Pfeffer et al., 2018). The time-varying amplitudes (power envelopes) were used to compute the inter-regional correlations in the MEG data (Fig. S2). Consequently, we eliminated all parameter combinations that fell outside of this regime of noise-driven oscillations from further consideration (see Methods for identification of the parameters producing sustained oscillations to be excluded).

##### Assumption 2. TASK increased the drive of both, E- and I-populations

We assumed that the change from REST to TASK corresponded to a shift of the model’s dynamical regime in an upward-rightward direction in the (***b_E_,b_I_***)-plane (Fig. 2C, area defined by dashed outline). This means that background input to both excitatory and inhibitory neural populations is increased for all nodes of the model. This assumption rests on the straightforward notion that our elementary visual task increased the input to sensory and task-related cortical regions. There is substantial evidence for the idea that cortical circuits generally operate in a regime of rough balance between excitation and inhibition (Shadlen and Newsome, 1998; van Vreeswijk and Sompolinsky, 1996). Specifically, sensory input increases not only feedforward excitation (i.e., feedforward drive of pyramidal cells), but also feedforward inhibition (i.e., feedforward drive of interneurons) in sensory cortex (McCasland and Hibbard, 1997; Swadlow, 2002), and it is assumed that this motif repeats across the cortical hierarchy (Shadlen and Newsome, 1998) likely augmented through circuit motifs for feedback inhibition (Womelsdorf et al., 2014). Correspondingly, we assumed that the visual task increased the background input to excitatory and inhibitory populations in a (loosely) balanced fashion, with a slight dominance of feedforward inhibition in the case of the cortex-wide model (see Fig. 2D,E). Indeed, recent evidence from rodent physiology shows that visual stimulation leads to a more pronounced inhibitory response (Adesnik, 2017; Haider et al., 2013) compared to the excitatory response, consistent with sensory input leading to even stronger feedforward inhibition compared to feedforward excitation. Note that this was in line with *Assumption 1*: if the task-induced increase in excitation was much larger than the task-induced increase in inhibition, the dynamical regime of the network would change to oscillatory, inconsistent with physiological evidence.

#### Simulation and fitting of cortex-wide Wilson Cowan model

The dynamical regime was defined as noise- or fluctuation-driven if: (1) the maximum and minimum were identical at any point in time or (2) the difference between maximum and minimum decreased monotonously over time (indicative of a damped oscillation). If none of the two were true, the signal was defined as a sustained oscillation. Note that this approach does not allow to distinguish between the two regimes with full certainty as the time scale with which the amplitude of an oscillation decays back to the fixed point increases as one approaches the Hopf-bifurcation from the fluctuation-driven regime (see Fig. S8A, middle and bottom panels). Thus, the closer the dynamical regime is to the Hopf-bifurcation, the more simulation time is required to accurately distinguish a sustained oscillation from a damped oscillation.

#### Microcircuit modeling (spiking neurons)

##### Microcircuit model of local node

We simulated the leaky integrate-and-fire circuit across a range of parameters to identify a stable working point where the network exhibits dynamics reminiscent of the “asynchronous state” (Renart et al., 2010). We defined the asynchronous state as being characterized by a low spontaneous firing rate (1-5 Hz) and low mean pairwise spike correlations (r < 0.1; averaged across all pairs of excitatory units). In addition, we identified a working point in the “synchronized state”, where the pairwise spike correlations where relatively high (r > 0.3), but spontaneous firing rates were comparable to the asynchronous state. This was achieved by changing AMPA-mediated recurrent excitation as well as the GABA-mediated feedback inhibition in a multiplicative manner: we started from the baseline parameters of Wang (2002), with some minor changes (see Methods), and multiplied *g_EE,AMPA_* (i.e., conductance of recurrent AMPA receptors) and *g_IE,GABA_* (i.e., conductance of GABAergic feedback inhibition) with 24 and 12 (respectively) linearly spaced values, ranging from 0.2 to 5 and 2.7 to 5. Fig. S10A shows spike rate (left) and mean pairwise spike correlations (right) for all parameter combinations. While for the main part of the analysis, we focus on the asynchronous state (shown in Fig. 2F; mean firing rate FR = 4.14 Hz; mean pairwise spike correlations r = 0.05), we further wanted to test whether the observed effect of feedback inhibition on response gain holds true also for the synchronous state. To this end, we first identified a dynamical regime reminiscent of the synchronous state, with low firing rates (FR = 5.11 Hz) but relatively high pairwise spike correlations (r = 0.27). We find that, irrespective of state, a reduction in feedback inhibition leads to an increase in response gain (Fig. S10B).

##### Tuning of parameters of decision circuit model

This model was based on a circuit model of decision-making developed to explain neural dynamics and choice behavior in standard two-alternative forced choice tasks, entailing trials of a few seconds of duration (Wang, 2002). Without further adjustments to the parameters of the decision circuit, the network dynamics would rapidly enter one of the two possible attractor states, reflecting the preference for decision 1 or decision 2. Moreover, without sufficient levels of external drive or noise, the network would dwell in those states indefinitely, as the lateral inhibition would dominate over the external input or the magnitude of the noise. In order to introduce dynamics that exceed beyond short timescales (single trials), we increased the level of background noise as well as the strength of the external stimulus. This way, we identified a state where the model would switch continuously between two attractors. Once this point was identified, we only changed feedback inhibition in order to assess the influence of E/I ratio on perceptual transitions (Fig. 3C).

Note that other studies that employed neural circuit models similar to the one used here for the study of perceptual fluctuations during ambiguous stimulations (Moreno-Bote et al., 2007) also included adaptation as an alternative mechanistic explanation for perceptual transitions. For the sake of simplicity, we did not consider this in the current circuit.

## Supplementary Figures

**Fig. S1.**
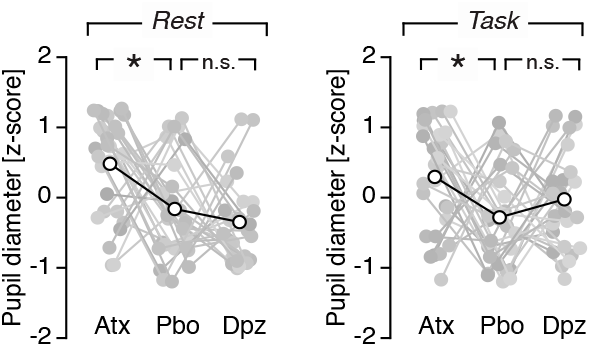
Drug effect on baseline pupil diameter. Baseline pupil diameter after the administration of atomoxetine (Atx), placebo (Pbo) or donepezil (Dpz), separately for rest (left) and task (right). (*) indicates *P* < 0.05 (paired t-test).

**Fig. S2.**
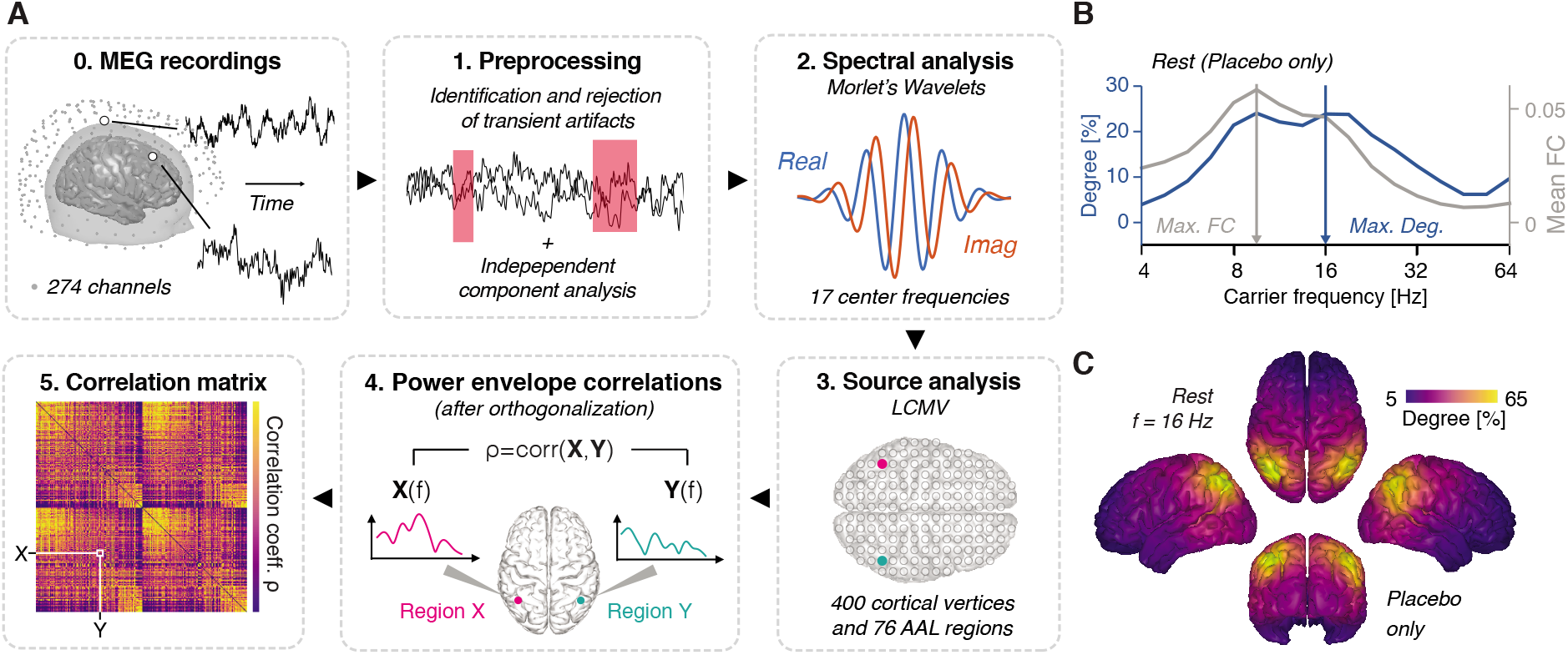
Quantifying cortex-wide correlation structure. **(A)** (0.) Whole-head magnetoencephalography (MEG) was recorded using 274 recording channels located in a helmet above the participants head. (**1**.) The sensor-level signal was cleaned from transient and sustained artifacts (e.g. muscle and heart beat artifacts, respectively). (2.) Spectral estimates were obtained from the cleaned sensor-level signal using complex wavelet convolution. (3.) From the spectral estimates and individual head models, source level power time series were obtained, (4.) from which orthogonalized power envelope correlations were computed. (5.) This resulted in functional connectivity (FC) matrices for each of the 17 carrier frequency bands of interest. **(B)** Global degree (see Methods) and mean FC as a function of frequency during the restplacebo condition. **(C)** Spatial map of degree during the rest-placebo condition. Correlations peak in the ‘alpha’ (center frequency 9.51 Hz; B) and ‘beta’ frequency range (center frequency 16 Hz; B), with strongest ‘connectedness’ (degree) in left and right posterior parietal cortex. These results are consistent with previous reports using an analogous approach to resting-state MEG data.

**Fig. S3.**
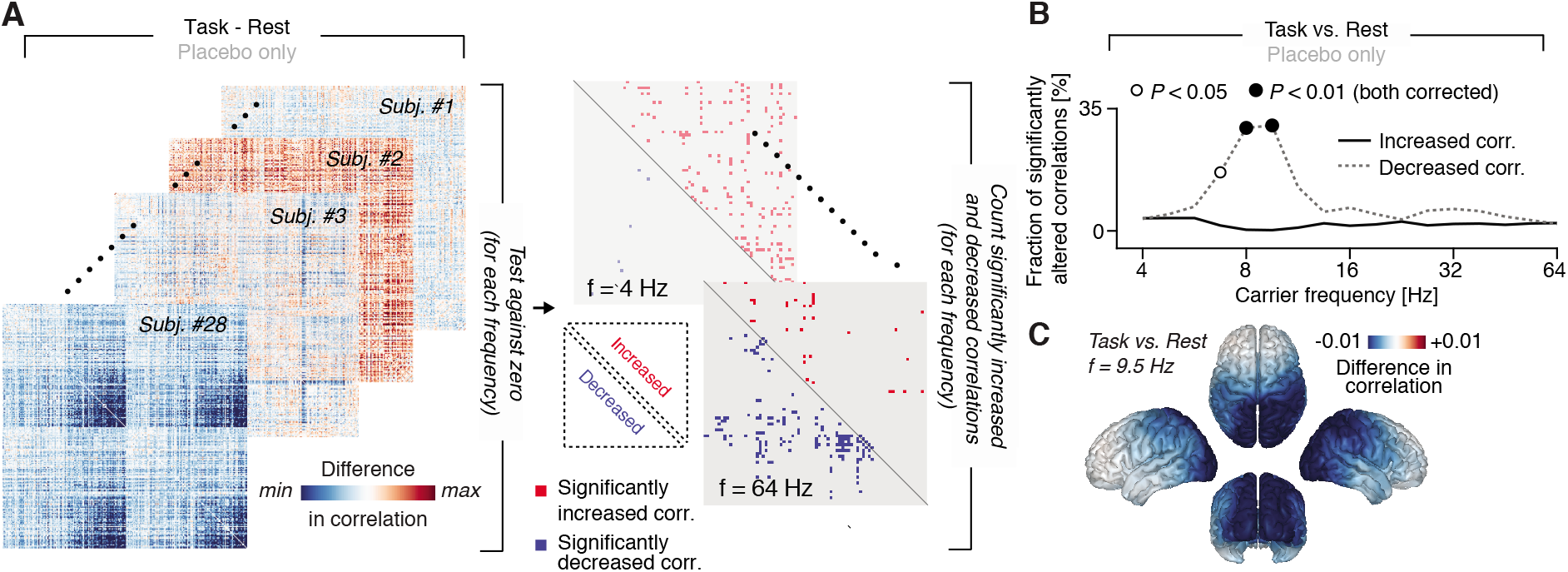
Quantifying task effects on cortical activity correlations. **(A)** Illustration of the approach to quantify changes in the global activity correlation structure: First, the difference between two conditions (here: rest and task; placebo only) was tested by means of a paired t-test. Next, the number of statistically significantly (*P* < 0.05; uncorrected) increased (in red) and significantly decreased (in blue) connections was counted. **(B)** A fraction of significantly altered correlations (negative and positive alterations) can be computed for each frequency band of interest, resulting in a spectrum of fraction of significantly altered correlations. **(C)** Spatial distribution of the difference in correlation between task and rest (placebo only) at a carrier frequency of 9.5 Hz.

**Fig. S4.**
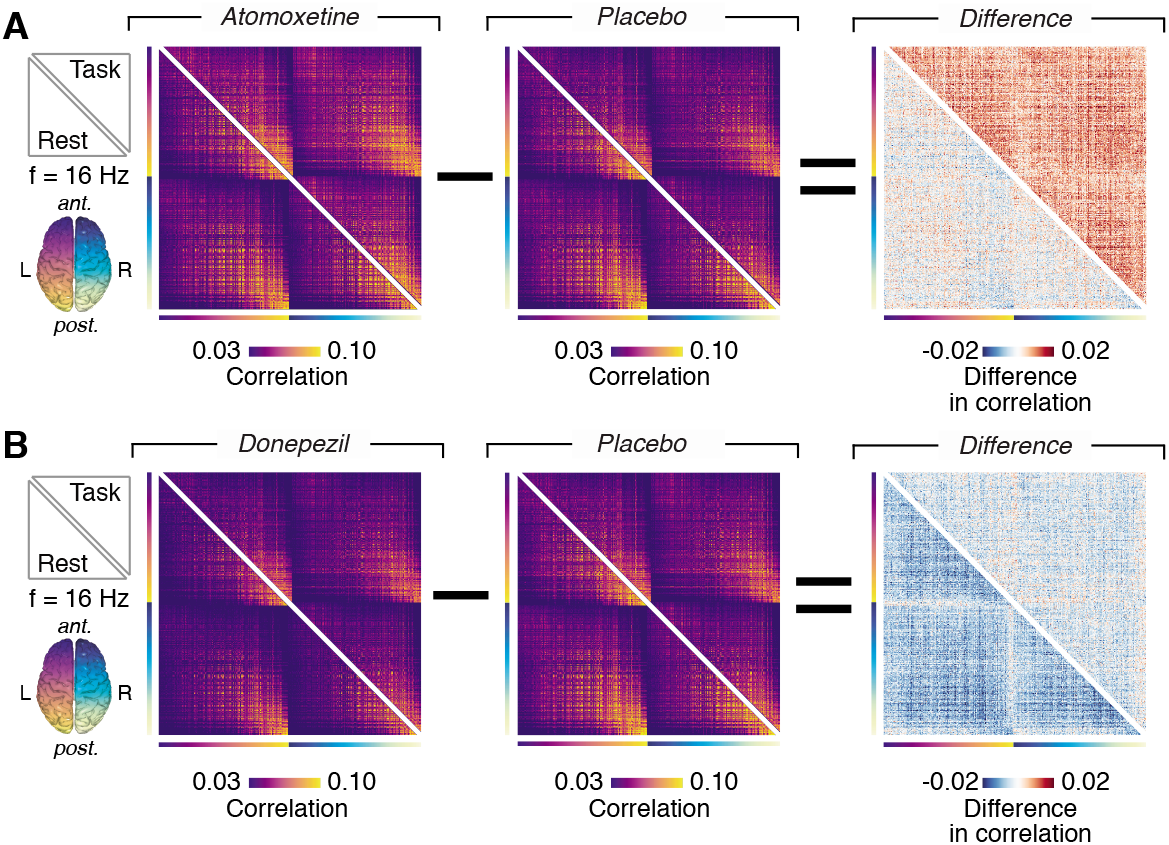
Quantifying drug effects on cortical activity correlations. **(A)** Functional connectivity matrices (only lower or upper triangular parts; at a carrier frequency of 16 Hz) for atomoxetine, placebo and the difference between the two (lower triangular part: during rest; upper triangular part: during task) and **(B)** for donepezil, placebo and the difference between the two.

**Fig. S5.**
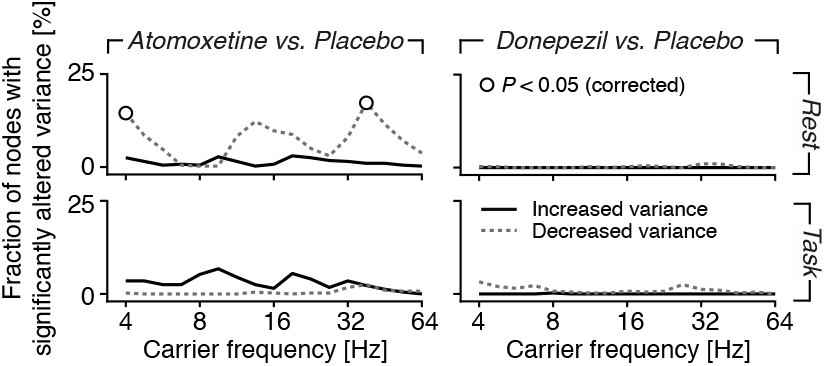
Changes in correlations were not driven by changes in local variance. Atomoxetine (left) induced a weak increase in the fraction of nodes with reduced power envelope variance compared to placebo during rest (open circles indicate *P* < 0.05, corrected for multiple comparisons across frequencies). Atomoxetine produced a tendency towards the opposite effect during task (increase in local variance) during task, albeit not statistically significant. Donepezil (right) did not lead to any significant alterations in local power envelope variance. The pattern of local variance changes under atomoxetine (decrease during rest, increase during task) cannot explain the observed pattern of changes in correlations under atomoxetine (no effect during rest, increase during task). In particular, an increase in local variance during task would reduce, not increase, correlations, because the local variance enters in the denominator in the computation of the correlation coefficient.

**Fig. S6.**
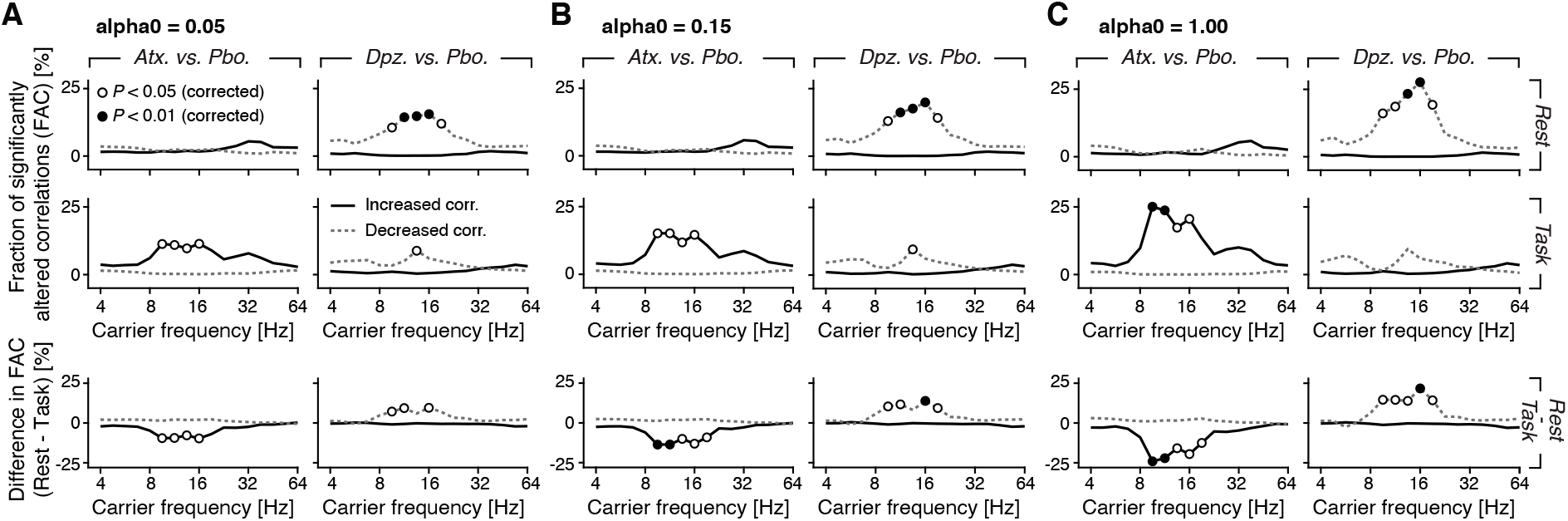
Fraction of significantly altered correlations for various regularization parameters. **(A-C)** Fraction of significantly altered correlations (FAC; as in Fig. 1E) for different regularization parameters used for the source reconstruction procedure (see Methods; panel A: α = 0.05; panel B: α = 0.15; panel C: α = 1.00). In the top row, the drug effects during rest are shown, in the middle row the effects during task (Left: Atomoxetine vs. placebo; Right: donepezil vs. placebo). The bottom row shows the effect of behavioral state (or context), i.e., the difference between the drug effect during Rest and the drug effect during Task. Significant differences are indicated by open circles (*P* < 0.05) and filled circles (*P* < 0.01; two-sided single-threshold permutation test).

**Fig. S7.**
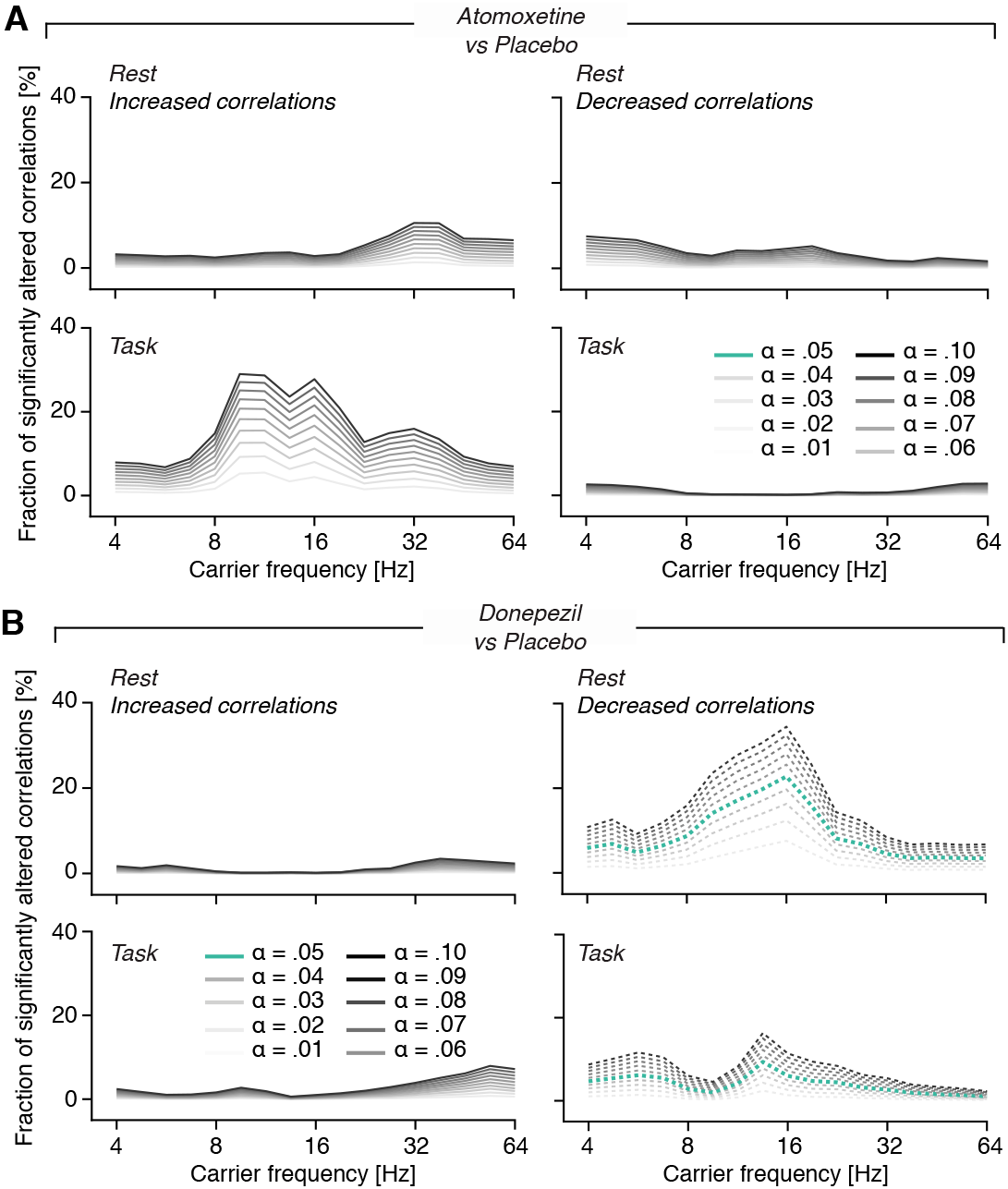
Fraction of significantly altered correlations for various alpha-thresholds parameters. The choice of the alpha-level for the initial paired t-test (Fig. 1E) does not affect the general qualitative pattern of the drug-induced changes in the fraction of significantly altered correlations. **(A)** Fraction of altered correlations as in Fig. 1E, but for various different alpha-values (for the initial t-test; see Methods), ranging from 0.01 to 0.10, for atomoxetine and **(B)** donepezil.

**Fig. S8.**
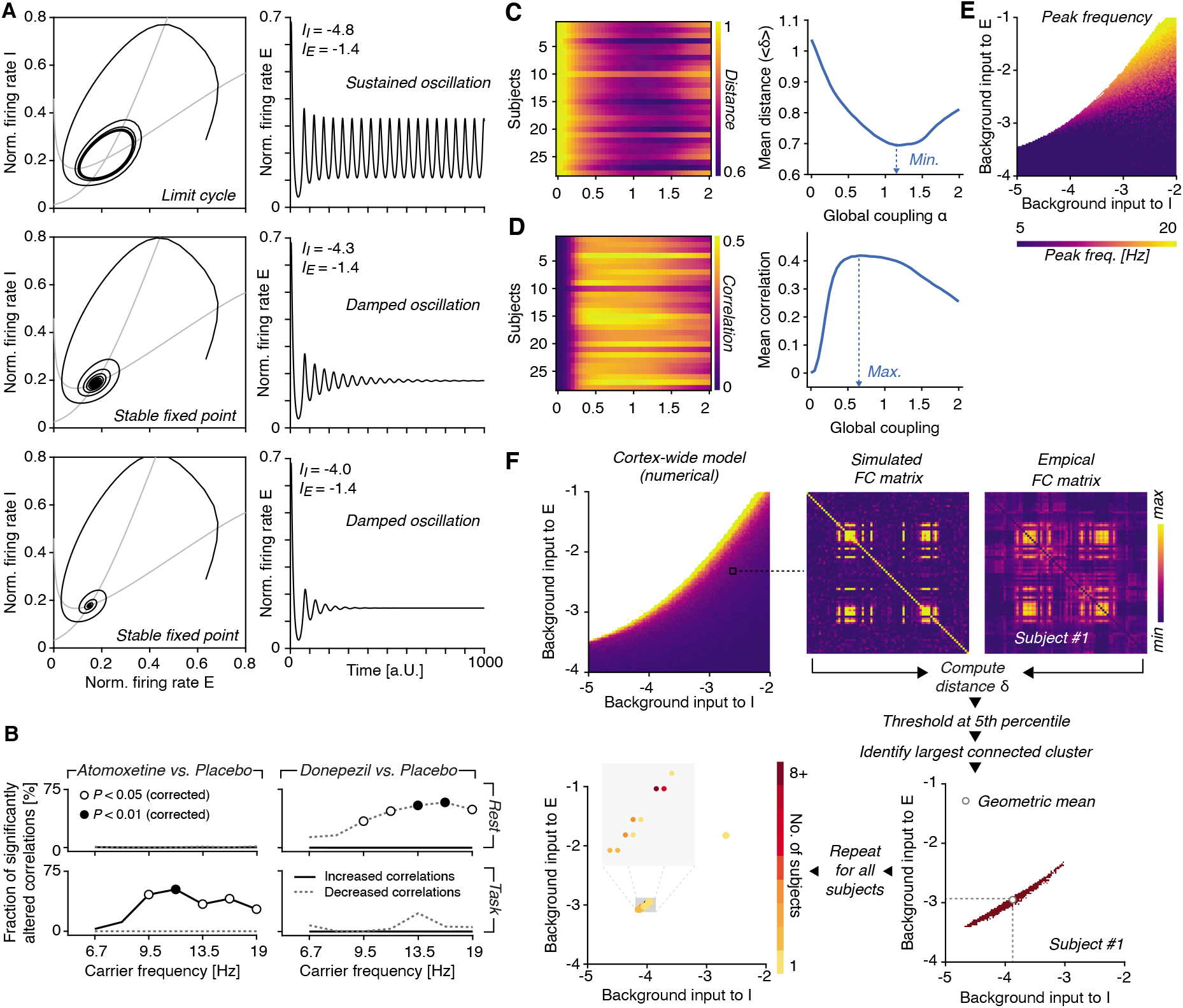
Dynamics of the Wilson-Cowan models and fitting procedure of the cortex-wide model. **(A)** Dynamics of a single Wilson-Cowan node around the Hopf-bifurcation. Top: Dynamics in the oscillatory regime, exhibiting sustained oscillations. Middle and bottom: Dynamics in the noise-driven regime, exhibiting damped oscillations. **(B)** The drug effects on fraction of significantly altered correlations for 76 AAL nodes and selected carrier frequency bands (ranging from ~6.7 to ~19 Hz). **(C)** Estimation of the global coupling parameter. Left: across all participants and a number of background input parameters (to E and I), the similarity (here: distance; see Methods) between the simulated and the empirical functional connectivity matrices was computed. Right: Distance (averaged across participants) for various levels of global coupling. **(D)** Same as (C), but for Pearson correlation. **(E)** Peak frequency of the model for various levels of background inputs to E and I. **(F)** Illustration of the fitting procedure: for each combination of background inputs to E and I, as well as every participant, the distance between simulated and empirical FC (rest and placebo only) was computed. The resulting distance matrix was thresholded at the 2.5^th^ percentile (all values larger were set to zero, all others to 1) and the largest connected cluster was identified. The geometric mean of this cluster was defined as the best-fitting value for a given participant. Repeating this procedure for all participants resulted in 28 fitted resting state parameters.

**Fig. S9.**
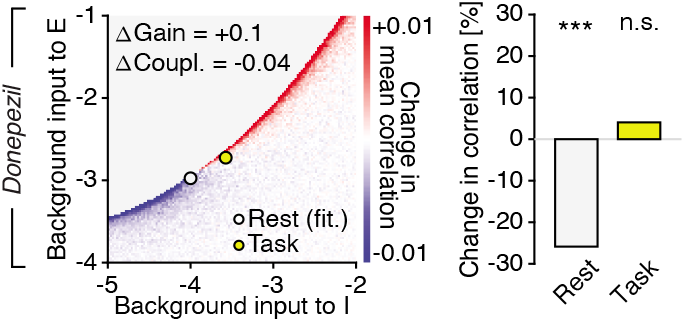
Effect of identical gain increase as in Fig. 2D (+0.1) combined with decrease of global coupling (−0.04) in cortex-wide model. This produces a similar decrease of cortex-wide correlations during rest as observed in the data under donepezil, but an increase in correlations during task, different from both, the donepezil and atomoxetine data. Note that the specific pattern of these effect depends on the amplitude of the task-related shift in the (*b_E_,* hy)**-**plane; the parameter changes used here may capture the observed donepezil effect with a different task-related shift in the (*b_E_,* Ďy)**-**plane. However, an equally big gain increase under acetylcholine as under catecholamines seems unlikely, given existing physiological data (Supplementary Information) and given the difference between the effects of both drugs on the number of perceptual transitions during the task (Fig. 3A and Fig. S11).

**Fig. S10.**
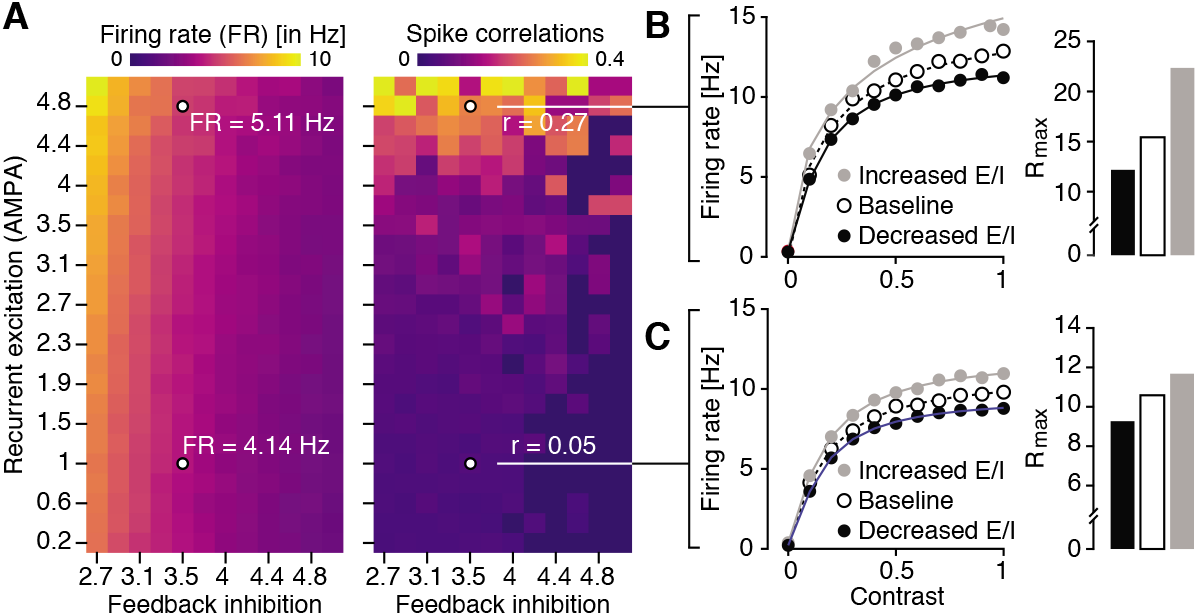
Increased E/I ratio increases response gain in asynchronous and synchronous states. **(A)** The parameters of the leaky integrate-and-fire model were tuned such that the network dynamics are indicative of a synchronized state (depicted pairwise spike count correlations r = 0.27) and low baseline firing rate (depicted FR = 5.11 Hz) or an asynchronous regime (pairwise spike count correlations r = 0.05), with comparable baseline firing rate (FR = 4.14 Hz). **(B)** The effects of altered feedback inhibition on response gain in the synchronous regime **(C)** Same as (B), but for the asynchronous regime (identical to Fig. 2F, replotted here for better comparison).

**Fig. S11.**
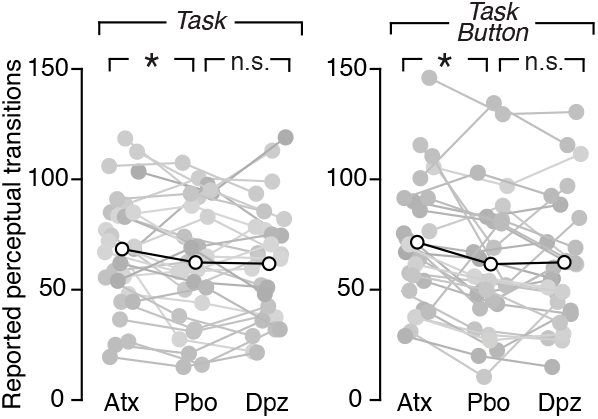
Behavior during task and task-pressing. Left: Number of reported perceptual transitions after the administration of atomoxetine (Atx), placebo (Pbo) or donepezil (Dpz), separately for task (silent counting of perceptual transitions). Right: same for task-button (perceptual transitions reported through pressing a button; right) (*) indicates *P* < 0.05 (two-sided paired t-test).

**Fig. S12.**
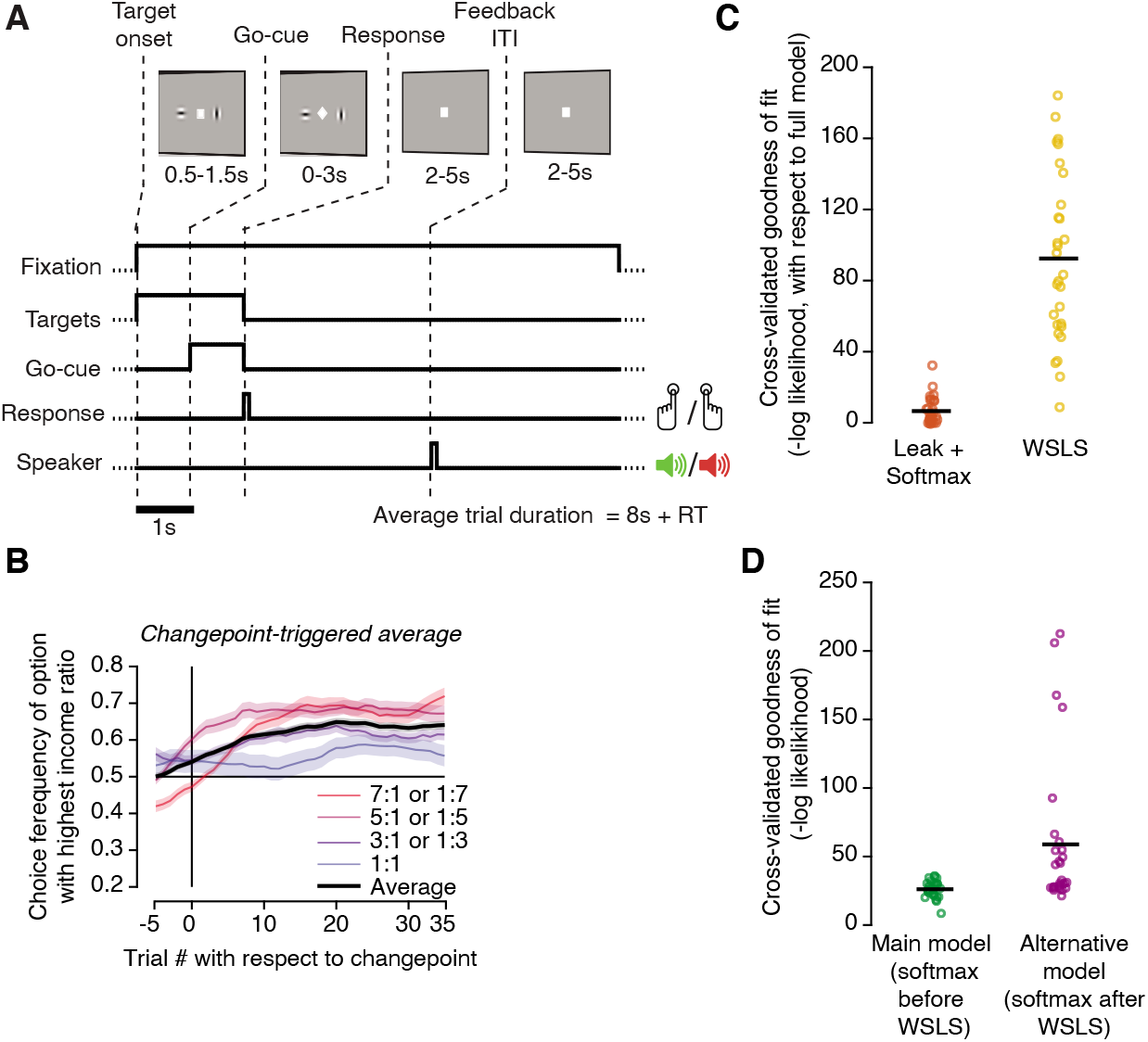
Task design, behavior, and behavioral modeling for dynamic foraging task. **(A)** Behavioral task. Top: sequence of events during each trial. Two choice targets (vertical/horizontal Gabors, randomized location) are presented at trial onset. A go-cue (change of fixation marker) instructs subjects to indicate their choice, by pressing a button with left or right index finger. Binary auditory feedback (reward or no-reward) is delivered after variable delay. **(B)** Change-point triggered change in choice fraction. **(C)** Cross-validated comparison between behavioral model from main Fig. 4F with a model entailing only WSLS or only leaky reward integration combined with softmax transformation. The latter fits the data better, indicating that a reward integration mechanism is needed to account for the data**. (D)** Cross-validated comparison between behavioral model from main Fig. 4F and a model, in which softmax transformation of choice probability is applied after combination with WSLS heuristic. The model from Fig. 4F (softmax transformation before WSLS) fits the data better.

**Table S1.** List of included AAL regions.

| REGION<br>INDEX | AAL<br>REGION NAME | REGION<br>INDEX | AAL<br>REGION NAME |
| --- | --- | --- | --- |
| 1 | Precentral_L | 39 | Temporal_Inf_R |
| 2 | Frontal_Sup_L | 40 | Temporal_Pole_Mid_R |
| 3 | Frontal_Sup_Orb_L | 41 | Temporal_Mid_R |
| 4 | Frontal_Mid_L | 42 | Temporal_Pole_Sup_R |
| 5 | Frontal_Mid_Orb_L | 43 | Temporal_Sub_R |
| 6 | Frontal_Inf_Oper_L | 44 | Heschl_R |
| 7 | Frontal_Inf_Tri_L | 45 | Paracentral_Lobule_R |
| 8 | Frontal_Inf_Orb_L | 46 | Precuneus_R |
| 9 | Rolandic_Oper_L | 47 | Angular_R |
| 10 | Supp_Motor_Area_L | 48 | SupraMarginal_R |
| 11 | Frontal_Supp_Medial_L | 49 | Parietal_Inf_R |
| 12 | Frontal_Med_Orb_L | 50 | Parietal_Sup_R |
| 13 | Rectus_L | 51 | Postcentral_R |
| 14 | Cingulum_Ant_L | 52 | Fusiform_R |
| 15 | Cingulum_Mid_L | 53 | Occipital_Inf_R |
| 16 | Cingulum_Post_L | 54 | Occipital_Mid_R |
| 17 | Hippocampus_L | 55 | Occipital_Sup_R |
| 18 | ParaHippocampal_L | 56 | Lingual_R |
| 19 | Calcarine_L | 57 | Cuneus_R |
| 20 | Cuneus_L | 58 | Calcarine_R |
| 21 | Lingual_L | 59 | ParaHippocampal_R |
| 22 | Occipital_Sup_L | 60 | Hippocampus_R |
| 23 | Occipital_Mid_L | 61 | Cingulum_Post_R |
| 24 | Occipital_Inf_L | 62 | Cingulum_Mid_R |
| 25 | Fusiform_L | 63 | Cingulum_Ant_R |
| 26 | Postcentral_L | 64 | Rectus_R |
| 27 | Parietal_Sup_L | 65 | Frontal_Med_Orb_R |
| 28 | Parietal_Inf_L | 66 | Frontal_Supp_Medial_R |
| 29 | SupraMarginal_L | 67 | Supp_Motor_Area__ |
| 30 | Angular_L | 68 | Rolandic_Oper_R |
| 31 | Precuneus_L | 69 | Frontal_Inf_Orb_R |
| 32 | Paracentral_Lobule_L | 70 | Frontal_Inf_Tri_R |
| 33 | Heschl_L | 71 | Frontal_Inf_Oper_R |
| 34 | Temporal_Sub_L | 72 | Frontal_Mid_Orb_R |
| 35 | Temporal_Pole_Sup_L | 73 | Frontal_Mid_R |
| 36 | Temporal_Mid_L | 74 | Frontal_Sup_Orb_R |
| 37 | Temporal_Pole_Mid_L | 75 | Frontal_Sup_R |
| 38 | Temporal_Inf_L | 76 | Precentral_R |

List of cortical AAL regions included in the model-based analysis.

## Movie S1. 3D-Structure-from-motion stimulus

https://www.youtube.com/watch?v=baZqACCbQqk

The 3D-Structure-from-Motion stimulus presented to the participants during the visual task.

## References

Adesnik, H. (2017). Synaptic Mechanisms of Feature Coding in the Visual Cortex of Awake Mice. Neuron 95, 1147–1159.e4.

Arnsten, A.F.T. (2015). Stress weakens prefrontal networks: molecular insults to higher cognition. Nat Neurosci 18, 1376–1385.

Arnsten, A.F.T., Paspalas, C.D., Gamo, N.J., Yang, Y., and Wang, M. (2010). Dynamic Network Connectivity: A new form of neuroplasticity. Trends Cogn. Sci. (Regul. Ed.) 14, 365–375.

Aston-Jones, G., and Cohen, J.D. (2005). An integrative theory of locus coeruleus-norepinephrine function: adaptive gain and optimal performance. Annu. Rev. Neurosci. 28, 403–450.

Aston-Jones, G., Rajkowski, J., and Cohen, J. (1999). Role of locus coeruleus in attention and behavioral flexibility. 46, 1309–1320.

Bear, M.F., and Singer, W. (1986). Modulation of visual cortical plasticity by acetylcholine and noradrenaline. Nature 320, 172–176.

Bell, A.J., and Sejnowski, T.J. (1995). An information-maximization approach to blind separation and blind deconvolution. Neural Comput 7, 1129–1159.

Brainard, D.H. (1997). The Psychophysics Toolbox. Spat Vis 10, 433–436.

Breton-Provencher, V., and Sur, M. (2019). Active control of arousal by a locus coeruleus GABAergic circuit. Nat Neurosci 22, 218–228.

van den Brink, R.L., Pfeffer, T., Warren, C.M., Murphy, P.R., Tona, K.-D., van der Wee, N.J.A., Giltay, E., van Noorden, M.S., Rombouts, S.A.R.B., Donner, T.H., et al. (2016). Catecholaminergic Neuromodulation Shapes Intrinsic MRI Functional Connectivity in the Human Brain. The Journal of Neuroscience 36, 7865–7876.

van den Brink, R.L., Nieuwenhuis, S., and Donner, T.H. (2018). Amplification and Suppression of Distinct Brainwide Activity Patterns by Catecholamines. J. Neurosci. 38, 7476–7491.

van den Brink, R.L., Pfeffer, T., and Donner, T.H. (2019). Brainstem Modulation of Large-Scale Intrinsic Cortical Activity Correlations. Front. Hum. Neurosci. 13, 340.

Brookes, M.J., Woolrich, M.W., and Barnes, G.R. (2012). Measuring functional connectivity in MEG: A multivariate approach insensitive to linear source leakage. NeuroImage 63, 910–920.

Burt, J.B., Demirtaş, M., Eckner, W.J., Navejar, N.M., Ji, J.L., Martin, W.J., Bernacchia, A., Anticevic, A., and Murray, J.D. (2018). Hierarchy of transcriptomic specialization across human cortex captured by structural neuroimaging topography. Nature Neuroscience.

Bymaster, F.P., Katner, J.S., Nelson, D.L., Hemrick-Luecke, S.K., Threlkeld, P.G., Heiligenstein, J.H., Morin, S.M., Gehlert, D.R., and Perry, K.W. (2002). Atomoxetine increases extracellular levels of norepinephrine and dopamine in prefrontal cortex of rat: a potential mechanism for efficacy in attention deficit/hyperactivity disorder. Neuropsychopharmacology 27, 699–711.

Castro-Alamancos, M.A., and Gulati, T. (2014). Neuromodulators produce distinct activated states in neocortex. J. Neurosci. 34, 12353–12367.

Cohen, J.D., McClure, S.M., and Yu, A.J. (2007). Should I stay or should I go? How the human brain manages the trade-off between exploitation and exploration. Phil. Trans. R. Soc. B 362, 933–942.

Cools, R. (2019). Chemistry of the Adaptive Mind: Lessons from Dopamine. Neuron 104, 113–131.

Corrado, G.S., Sugrue, L.P., Sebastian Seung, H., and Newsome, W.T. (2005). Linear-Nonlinear-Poisson Models of Primate Choice Dynamics. Journal of the Experimental Analysis of Behavior 84, 581–617.

Coull, J.T., Büchel, C., Friston, K.J., and Frith, C.D. (1999). Noradrenergically Mediated Plasticity in a Human Attentional Neuronal Network. NeuroImage 10, 705–715.

Deco, G., Jirsa, V., McIntosh, A.R., Sporns, O., and Kotter, R. (2009). Key role of coupling, delay, and noise in resting brain fluctuations. Proceedings of the National Academy of Sciences 106, 10302–10307.

Deco, G., Ponce-Alvarez, A., Hagmann, P., Romani, G.L., Mantini, D., and Corbetta, M. (2014). How local excitation-inhibition ratio impacts the whole brain dynamics. J. Neurosci. 34, 7886–7898.

Deco, G., Kringelbach, M.L., Jirsa, V.K., and Ritter, P. (2017). The dynamics of resting fluctuations in the brain: metastability and its dynamical cortical core. Sci Rep 7, 3095.

Deco, G., Cruzat, J., Cabral, J., Knudsen, G.M., Carhart-Harris, R.L., Whybrow, P.C., Logothetis, N.K., and Kringelbach, M.L. (2018). Whole-Brain Multimodal Neuroimaging Model Using Serotonin Receptor Maps Explains Non-linear Functional Effects of LSD. Current Biology 28, 3065–3074.e6.

Demirtaş, M., Burt, J.B., Helmer, M., Ji, J.L., Adkinson, B.D., Glasser, M.F., Van Essen, D.C., Sotiropoulos, S.N., Anticevic, A., and Murray, J.D. (2019a). Hierarchical Heterogeneity across Human Cortex Shapes Large-Scale Neural Dynamics. Neuron 101, 1181–1194.e13.

Demirtaş, M., Burt, J.B., Helmer, M., Ji, J.L., Adkinson, B.D., Glasser, M.F., Van Essen, D.C., Sotiropoulos, S.N., Anticevic, A., and Murray, J.D. (2019b). Hierarchical Heterogeneity across Human Cortex Shapes Large-Scale Neural Dynamics. Neuron 101, 1181–1194.e13.

Disney, A. A., Aoki, C., and Hawken, M.J. (2007). Gain Modulation by Nicotine in Macaque V1. Neuron 56, 701–713.

Eldar, E., Cohen, J.D., and Niv, Y. (2013). The effects of neural gain on attention and learning. Nat Neurosci 16, 1146–1153.

Ferguson, K.A., and Cardin, J.A. (2020). Mechanisms underlying gain modulation in the cortex. Nat Rev Neurosci 21, 80–92.

Findling, C., Skvortsova, V., Dromnelle, R., Palminteri, S., and Wyart, V. (2019). Computational noise in reward-guided learning drives behavioral variability in volatile environments. Nat Neurosci 22, 2066–2077.

Fox, M.D., and Greicius, M. (2010). Clinical applications of resting state functional connectivity. Front. Syst. Neurosci.

Frank, M.J., Doll, B.B., Oas-Terpstra, J., and Moreno, F. (2009). Prefrontal and striatal dopaminergic genes predict individual differences in exploration and exploitation. Nature Neuroscience 12, 1062–1068.

Froemke, R.C. (2015). Plasticity of Cortical Excitatory-Inhibitory Balance. Annual Review of Neuroscience 38, 195–219.

Froemke, R.C., Merzenich, M.M., and Schreiner, C.E. (2007). A synaptic memory trace for cortical receptive field plasticity. Nature 450, 425–429.

de Gee, J.W., Colizoli, O., Kloosterman, N.A., Knapen, T., Nieuwenhuis, S., and Donner, T.H. (2017). Dynamic modulation of decision biases by brainstem arousal systems. ELife 6.

Goodman, D.F.M. (2009). The Brian simulator. Frontiers in Neuroscience 3, 192–197.

Goodman, D.F., Stimberg, M., Yger, P., and Brette, R. (2014). Brian 2: neural simulations on a variety of computational hardware. BMC Neuroscience 15, P199.

Gratton, C., Yousef, S., Aarts, E., Wallace, D.L., D’Esposito, M., and Silver, M.A. (2017). Cholinergic, But Not Dopaminergic or Noradrenergic, Enhancement Sharpens Visual Spatial Perception in Humans. J. Neurosci. 37, 4405–4415.

Haider, B., Häusser, M., and Carandini, M. (2013). Inhibition dominates sensory responses in the awake cortex. Nature 493, 97–100.

Harris, K.D., and Thiele, A. (2011). Cortical state and attention. Nat. Rev. Neurosci. 12, 509–523.

Hawellek, D.J., Schepers, I.M., Roeder, B., Engel, A.K., Siegel, M., and Hipp, J.F. (2013). Altered Intrinsic Neuronal Interactions in the Visual Cortex of the Blind. Journal of Neuroscience 33, 17072–17080.

Herrero, J.L., Roberts, M.J., Delicato, L.S., Gieselmann, M.A., Dayan, P., and Thiele, A. (2008). Acetylcholine contributes through muscarinic receptors to attentional modulation in V1. Nature 454, 1110–1114.

Herrero, J.L., Gieselmann, M.A., and Thiele, A. (2017). Muscarinic and Nicotinic Contribution to Contrast Sensitivity of Macaque Area V1 Neurons. Front. Neural Circuits 11, 106.

Hipp, J.F., and Siegel, M. (2013). Dissociating neuronal gamma-band activity from cranial and ocular muscle activity in EEG. Frontiers in Human Neuroscience 7.

Hipp, J.F., Hawellek, D.J., Corbetta, M., Siegel, M., and Engel, A.K. (2012). Large-scale cortical correlation structure of spontaneous oscillatory activity. Nature Neuroscience 15, 884–890.

Hsieh, C.Y., Cruikshank, S.J., and Metherate, R. (2000). Differential modulation of auditory thalamocortical and intracortical synaptic transmission by cholinergic agonist. Brain Res. 880, 51–64.

Hurley, L.M., Devilbiss, D.M., and Waterhouse, B.D. (2004). A matter of focus: monoaminergic modulation of stimulus coding in mammalian sensory networks. Curr. Opin. Neurobiol. 14, 488–495.

Hyvarinen, A. (1999). Fast and robust fixed-point algorithms for independent component analysis. IEEE Transactions on Neural Networks 10, 626–634.

Joshi, S., Li, Y., Kalwani, R.M., and Gold, J.I. (2016). Relationships between Pupil Diameter and Neuronal Activity in the Locus Coeruleus, Colliculi, and Cingulate Cortex. Neuron 89, 221–234.

Kane, G.A., Vazey, E.M., Wilson, R.C., Shenhav, A., Daw, N.D., Aston-Jones, G., and Cohen, J.D. (2017). Increased locus coeruleus tonic activity causes disengagement from a patch-foraging task. Cogn Affect Behav Neurosci 17, 1073–1083.

Knapen, T., de Gee, J.W., Brascamp, J., Nuiten, S., Hoppenbrouwers, S., and Theeuwes, J. (2016). Cognitive and Ocular Factors Jointly Determine Pupil Responses under Equiluminance. PLoS ONE 11, e0155574.

Lam, N.H., Borduqui, T., Hallak, J., Roque, A.C., Anticevic, A., Krystal, J.H., Wang, X.-J., and Murray, J.D. (2017). Effects of Altered Excitation-Inhibition Balance on Decision Making in a Cortical Circuit Model. BioRxiv.

Leopold, D., and Logothetis, N. (1999). Multistable phenomena: changing views in perception. 3, 254–264.

Leopold, D.A., Murayama, Y., and Logothetis, N.K. (2003). Very slow activity fluctuations in monkey visual cortex: implications for functional brain imaging. Cereb. Cortex 13, 422–433.

Letzkus, J.J., Wolff, S.B.E., Meyer, E.M.M., Tovote, P., Courtin, J., Herry, C., and Lüthi, A. (2011). A disinhibitory microcircuit for associative fear learning in the auditory cortex. Nature 480, 331–335.

Martins, A.R.O., and Froemke, R.C. (2015). Coordinated forms of noradrenergic plasticity in the locus coeruleus and primary auditory cortex. Nature Neuroscience 18, 1483–1492.

McCasland, J.S., and Hibbard, L.S. (1997). GABAergic Neurons in Barrel Cortex Show Strong, Whisker-Dependent Metabolic Activation during Normal Behavior. J. Neurosci. 17, 5509–5527.

McGinley, M.J., David, S.V., and McCormick, D.A. (2015). Cortical Membrane Potential Signature of Optimal States for Sensory Signal Detection. Neuron 87, 179–192.

Montague, P.R., Hyman, S.E., and Cohen, J.D. (2004). Computational roles for dopamine in behavioural control. Nature 431, 760–767.

Murphy, B.K., and Miller, K.D. (2003). Multiplicative gain changes are induced by excitation or inhibition alone. J. Neurosci. 23, 10040–10051.

Nichols, T.E., and Holmes, A.P. (2002). Nonparametric permutation tests for functional neuroimaging: A primer with examples. Human Brain Mapping 15, 1–25.

Oostenveld, R., Fries, P., Maris, E., and Schoffelen, J.-M. (2011). FieldTrip: Open source software for advanced analysis of MEG, EEG, and invasive electrophysiological data. Comput Intell Neurosci 2011, 156869.

Pfeffer, T., Avramiea, A.-E., Nolte, G., Engel, A.K., Linkenkaer-Hansen, K., and Donner, T.H. (2018). Catecholamines alter the intrinsic variability of cortical population activity and perception. PLOS Biology 16, e2003453.

Polack, P.-O., Friedman, J., and Golshani, P. (2013). Cellular mechanisms of brain state-dependent gain modulation in visual cortex. Nat. Neurosci. 16, 1331–1339.

Reimer, J., McGinley, M.J., Liu, Y., Rodenkirch, C., Wang, Q., McCormick, D.A., and Tolias, A.S. (2016). Pupil fluctuations track rapid changes in adrenergic and cholinergic activity in cortex. Nature Communications 7, 13289.

Renart, A., and Machens, C.K. (2014). Variability in neural activity and behavior. Curr. Opin. Neurobiol. 25, 211–220.

Robbins, T.W., and Arnsten, A.F.T. (2009). The neuropsychopharmacology of fronto-executive function: monoaminergic modulation. Annu. Rev. Neurosci. 32, 267–287.

Roberts, M.J., Zinke, W., Guo, K., Robertson, R., McDonald, J.S., and Thiele, A. (2005). Acetylcholine dynamically controls spatial integration in marmoset primary visual cortex. J. Neurophysiol. 93, 2062–2072.

Rubinov, M., and Sporns, O. (2010). Complex network measures of brain connectivity: uses and interpretations. Neuroimage 52, 1059–1069.

Sauer, J.-M., Ring, B.J., and Witcher, J.W. (2005). Clinical pharmacokinetics of atomoxetine. Clin Pharmacokinet 44, 571–590.

Schwarz, L.A., and Luo, L. (2015). Organization of the Locus Coeruleus-Norepinephrine System. Current Biology 25, R1051–R1056.

Servan-Schreiber, D., Printz, H., and Cohen, J. (1990). A network model of catecholamine effects: gain, signal-to-noise ratio, and behavior. Science 249, 892–895.

Shine, J.M., Aburn, M.J., Breakspear, M., and Poldrack, R.A. (2018). The modulation of neural gain facilitates a transition between functional segregation and integration in the brain. ELife 7.

Siems, M., Pape, A., Hipp, J., and Siegel, M. (2016). Measuring the cortical correlation structure of spontaneous oscillatory activity with EEG and MEG. NeuroImage.

Silver, M.A., Shenhav, A., and D’Esposito, M. (2008). Cholinergic enhancement reduces spatial spread of visual responses in human early visual cortex. Neuron 60, 904–914.

Soma, S., Shimegi, S., Osaki, H., and Sato, H. (2012). Cholinergic modulation of response gain in the primary visual cortex of the macaque. J. Neurophysiol. 107, 283–291.

Sugrue, L.P., Corrado, G.S., and Newsome, W.T. (2004). Matching behavior and the representation of value in the parietal cortex. Science 304, 1782–1787.

Sugrue, L.P., Corrado, G.S., and Newsome, W.T. (2005). Choosing the greater of two goods: neural currencies for valuation and decision making. Nat Rev Neurosci 6, 363–375.

Swadlow, H.A. (2002). Thalamocortical control of feed-forward inhibition in awake somatosensory “barrel” cortex. Philos. Trans. R. Soc. Lond., B, Biol. Sci. 357, 1717–1727.

Tallon-Baudry, C, and Bertrand, O (1999). Oscillatory gamma activity in humans and its role in object representation. Trends Cogn. Sci. (Regul. Ed.) 3, 151–162.

Tervo, D.G.R., Proskurin, M., Manakov, M., Kabra, M., Vollmer, A., Branson, K., and Karpova, A.Y. (2014). Behavioral Variability through Stochastic Choice and Its Gating by Anterior Cingulate Cortex. Cell 159, 21–32.

Tiseo, Rogers, and Friedhoff (1998). Pharmacokinetic and pharmacodynamic profile of donepezil HCl following evening administration: Evening administration of donepezil HCl. British Journal of Clinical Pharmacology 46, 13–18.

Turchi, J., Chang, C., Ye, F.Q., Russ, B.E., Yu, D.K., Cortes, C.R., Monosov, I.E., Duyn, J.H., and Leopold, D.A. (2018). The Basal Forebrain Regulates Global Resting-State fMRI Fluctuations. Neuron 97, 940–952.e4.

Tzourio-Mazoyer, N., Landeau, B., Papathanassiou, D., Crivello, F., Etard, O., Delcroix, N., Mazoyer, B., and Joliot, M. (2002). Automated Anatomical Labeling of Activations in SPM Using a Macroscopic Anatomical Parcellation of the MNI MRI Single-Subject Brain. NeuroImage 15, 273–289.

Usher, M., Cohen, J.D., Servan-Schreiber, D., Rajkowski, J., and Aston-Jones, G. (1999). The role of locus coeruleus in the regulation of cognitive performance. Science 283, 549–554.

Veen, V.B., van Drongelen, W., Yuchtman, M., and Suzuki, A. (1997). Localization of brain electrical activity via linearly constrained minimum variance spatial filtering. Biomedical … 44, 867–880.

Wallach, H., and O’connell, D.N. (1953). The kinetic depth effect. J Exp Psychol 45, 205–217.

Wang, X.-J. (2002). Probabilistic decision making by slow reverberation in cortical circuits. Neuron 36, 955–968.

Warren, C.M., Wilson, R.C., van der Wee, N.J., Giltay, E.J., van Noorden, M.S., Cohen, J.D., and Nieuwenhuis, S. (2017). The effect of atomoxetine on random and directed exploration in humans. PLoS ONE 12, e0176034.

Wilson, H.R., and Cowan, J.D. (1972). Excitatory and Inhibitory Interactions in Localized Populations of Model Neurons. Biophysical Journal 12, 1–24.

Wimmer, K., Compte, A., Roxin, A., Peixoto, D., Renart, A., and de la Rocha, J. (2015). Sensory integration dynamics in a hierarchical network explains choice probabilities in cortical area MT. Nature Communications 6.

Yu, A.J., and Dayan, P. (2005). Uncertainty, neuromodulation, and attention. Neuron 46, 681–692.

## Supplementary References

Beggs, J.M., and Plenz, D. (2003). Neuronal avalanches in neocortical circuits. J. Neurosci. 23, 11167–11177.

Moreno-Bote, R., Rinzel, J., and Rubin, N. (2007). Noise-induced alternations in an attractor network model of perceptual bistability. J. Neurophysiol. 98, 1125–1139.

Poil, S.-S., Hardstone, R., Mansvelder, H.D., and Linkenkaer-Hansen, K. (2012). Critical-State Dynamics of Avalanches and Oscillations Jointly Emerge from Balanced Excitation/Inhibition in Neuronal Networks. Journal of Neuroscience 32, 9817–9823.

Renart, A., Rocha, J., Bartho, P., Hollender, L., Parga, N., Reyes, A., and Harris, K.D. (2010). The asynchronous state in cortical circuits. 327, 587–590.

Shadlen, M.N., and Newsome, W.T. (1998). The variable discharge of cortical neurons: implications for connectivity, computation, and information coding. J. Neurosci. 18, 3870–3896.

van Vreeswijk, C., and Sompolinsky, H. (1996). Chaos in neuronal networks with balanced excitatory and inhibitory activity. Science 274, 1724–1726.

Womelsdorf, T., Valiante, T.A., Sahin, N.T., Miller, K.J., and Tiesinga, P. (2014). Dynamic circuit motifs underlying rhythmic gain control, gating and integration. Nat Neurosci 17, 1031–1039.

